# CTCF-dependent chromatin boundaries formed by asymmetric nucleosome arrays with decreased linker length

**DOI:** 10.1101/618827

**Authors:** Christopher T. Clarkson, Emma A. Deeks, Ralph Samarista, Hulkar Mamayusupova, Victor B. Zhurkin, Vladimir B. Teif

**Author notes:** Correspondence should be addressed to Vladimir B. Teif.

## Abstract

The CCCTC-binding factor (CTCF) organises the genome in 3D through DNA loops and in 1D by setting boundaries isolating different chromatin states, but these processes are not well understood. Here we focus on the relationship between CTCF binding and the decrease of the Nucleosome Repeat Length (NRL) for ∼20 adjacent nucleosomes, affecting up to 10% of the mouse genome. We found that the chromatin boundary near CTCF is created by the nucleosome-depleted region (NDR) asymmetrically located >40 nucleotides 5’-upstream from the centre of CTCF motif. The strength of CTCF binding to DNA is correlated with the decrease of NRL near CTCF and anti-correlated with the level of asymmetry of the nucleosome array. Individual chromatin remodellers have different contributions, with Snf2h having the strongest effect on the NRL decrease near CTCF and Chd4 playing a major role in the symmetry breaking. Upon differentiation of embryonic stem cells to neural progenitor cells and embryonic fibroblasts, a subset of common CTCF sites preserved in all three cell types maintains a relatively small local NRL despite genome-wide NRL increase. The sites which lost CTCF upon differentiation are characterised by nucleosome rearrangement 3’-downstream, but the boundary defined by the NDR 5’-upstream of CTCF motif remains.

## Introduction

Nucleosomes are positioned along the genome in a non-random way (1–3), which is critical for determining the DNA accessibility and genome organisation (4). A particularly important nucleosome positioning signal is provided by CTCF, an architectural protein that maintains 3D genome architecture (5–7) and can organise up to 20 nucleosomes in its vicinity (8) (Figure 1A). CTCF has ∼100,000 potential binding sites in the mouse genome. Usually there are ∼30,000-60,000 CTCF sites bound in a given cell type, which translates to about 1 million of affected nucleosomes (up to 10% of the mouse genome) (9–12). CTCF is able to act as an insulator between genomic regions with different chromatin states, but how exactly this is achieved is not known. Here we explore molecular mechanisms of the insulator boundary formation by CTCF through rearrangement of surrounding nucleosome arrays.

**Figure 1.**
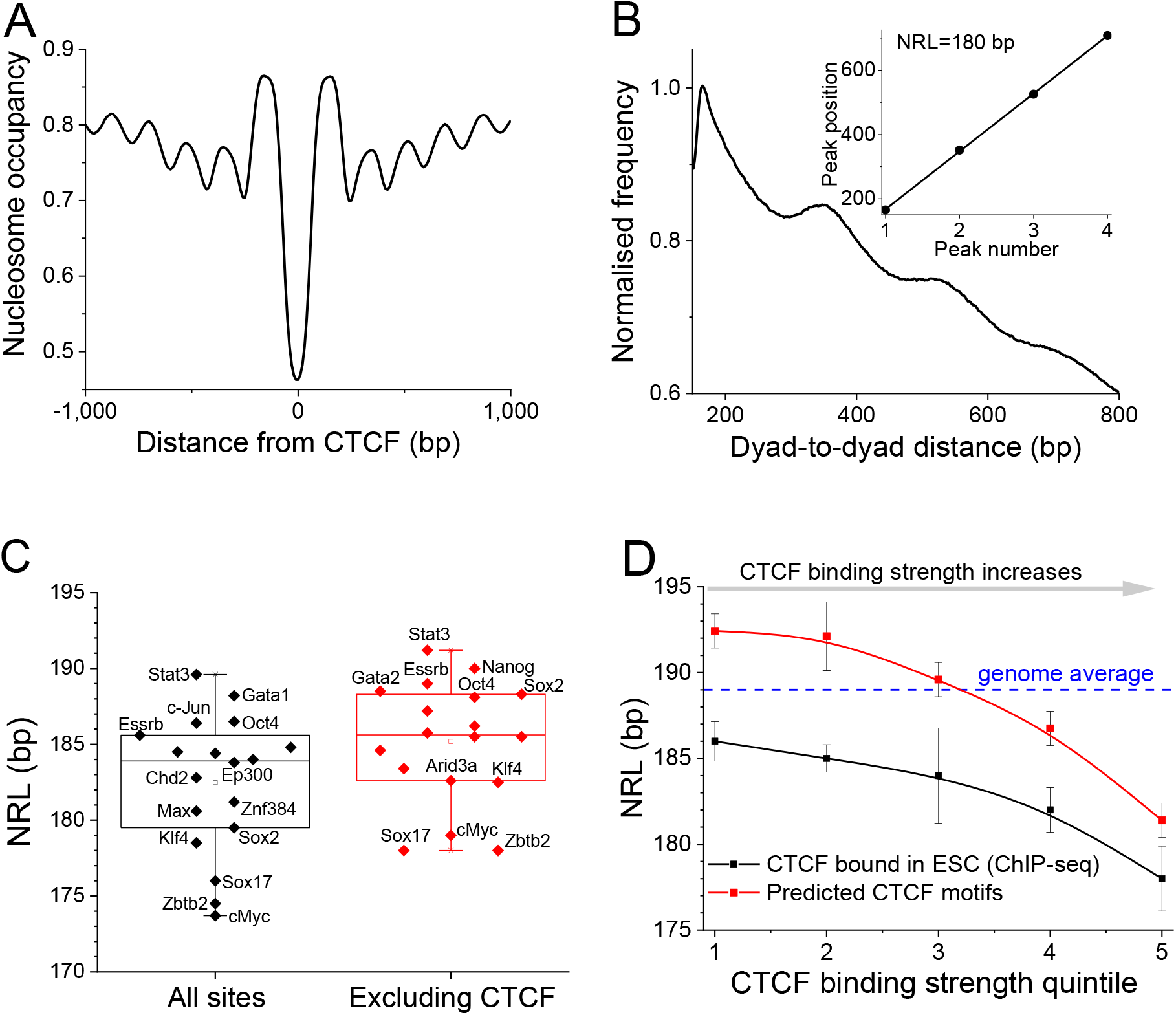
CTCF-dependent decrease of the nucleosome repeat length (NRL). A) Average nucleosome profile around CTCF binding sites in ESCs. B) The illustration of the “phasogram” method of NRL calculation for the region [100, 2000] from the centre of experimental CTCF sites measured in ESCs. The calculation of frequencies of nucleosome dyad-to-dyad distances is followed by the linear regression of the peak positions (insert). C) NRLs calculated near binding sites of 18 stemness-related chromatin proteins in ESCs in the region [100, 2000] from the summit of TF binding ChIP-seq peak, using chemical nucleosome mapping data from Voong et al (41). Left: all TF binding sites; right: TF binding sites which do not intersect with CTCF. D) Dependence of NRL on the strength of CTCF binding based on experimental ChIP-seq peaks from mouse ENCODE (9) stratified into binding strength quintiles by the heights of peaks (black line) and computationally predicted CTCF sites obtained by scanning the mouse genome with TFBStools using >80% similarity for JASPAR matrix MA0139.1 stratified into binding strength quintiles by their TRAP score (red line).

One of the ways to characterise genomic nucleosome distribution is through an integral parameter called the nucleosome repeat length (NRL), defined as the average distance between the centres of adjacent nucleosomes. NRL can be defined genome-wide, locally for an individual genomic region or for a set of regions. The local NRL is particularly important, since it reflects different structures of chromatin fibers (13–17). Ever since the discovery of the nucleosome (18,19) there have been many attempts to compare NRLs of different genomic regions (20–22) and it has been established that genome-wide NRL changes during cell differentiation (23,24). Recent sequencing-based investigations showed that active regions such as promoters, enhancers and actively transcribed genes usually have shorter NRLs while heterochromatin is characterised by longer NRLs (25–28). While in Yeast it is possible to link NRL changes to the action of individual chromatin remodellers (29–33), in higher eukaryotes regulatory regions are very heterogeneous and it is difficult to come up with a set of definitive remodeller rules determining their effect on NRL (34,35).

We previously showed that in mouse embryonic stem cells (ESC), NRL near CTCF is about 10 bp smaller than genome-wide NRL (36,37). Our analysis demonstrated that purely statistical positioning of nucleosomes near CTCF boundaries would result in a longer NRL than observed experimentally, and the effects of strong nucleosome-positioning DNA sequences, while compatible with the observed NRL, are limited to a small number of CTCF sites (38). A very recent study has investigated the effect of Snf2 and Brg1 remodellers on NRL in ESCs, suggesting Snf2 as the primary player (39). However, other factors may be at play as well. Thus, it is still unclear what determines the NRL near CTCF and how different CTCF sites are distinguished from each other e.g. during cell differentiation. Furthermore, recent studies have shown that CTCF can act as a boundary element between different chromatin states (e.g. DNA methylation) linearly spreading along the genome (10,40), but the mechanistic explanation for such a function is not immediately clear from the better established role of CTCF in 3D chromatin looping. Here we address these problems using available experimental datasets in ESCs and their differentiated counterparts.

We show below that the boundaries of nucleosome arrays are encoded in extended DNA regions >200 bp long enclosing the CTCF motifs. Furthermore, the strength of CTCF binding provides a single “code” that determines the value of NRL near CTCF, the level of asymmetry of CTCF-dependent nucleosome array boundaries, and eventually serves as a guide for chromatin rearrangements during cell differentiation.

## Materials and Methods

### Experimental datasets

Nucleosome positioning and transcription factor binding datasets were obtained from the Gene Expression Omnibus (GEO), Short Read Archive (SRA) and the ENCODE web site as detailed in Table ST1. NRL calculations near CTCF in ESCs were performed using the MNase-seq dataset from (41). NRL calculations near 19 stemness-related proteins in ESCs shown in Figure 1D and S1 were performed using the chemical mapping dataset from (41). NRL calculations in NPCs and MEFs were based on the MNase-seq datasets from (36). MNase-assisted H3 ChIP-seq from (10) was used for demonstrative purposes in the phasogram calculation in Figure 1C. Coordinates of genomic features and experimental maps of transcription factor and remodeller binding in ESCs were obtained from published sources as detailed in Table S1. The coordinates of loops and TADs described in (42) were provided by the authors in a BED file aligned to the mm10 mouse genome and were converted to mm9 using liftOver (UCSC Genome Browser).

### Data pre-processing

For nucleosome positioning, raw sequencing data were aligned to the mouse mm9 genome using Bowtie allowing up to 2 mismatches. For all other datasets we used processed files with genomic coordinates downloaded from the corresponding database as detailed in Table ST1. Where required, coordinates were converted from mm10 to mm9 since the majority of the datasets were in mm9.

### Basic data processing

TF binding-sites were extended from the center of the site to the region [100, 2000]. In order to find all nucleosomal DNA fragments inside each genomic region of interest the bed files containing the coordinates of nucleosomes processed using the NucTools pipeline (43) were intersected with the corresponding genomic regions of interest using BEDTools (44). Average nucleosome occupancy profiles were calculated using NucTols. The phasograms were calculated using NucTools as detailed below.

### Binding site prediction

Computationally predicted TF binding sites were determined via scanning the mouse genome with position frequency matrices (PFMs) from the JASPAR2018 database (45) using R packages TFBSTools (46) and GenomicRanges (47). A similarity threshold of 80% was used for all TFs in order to get at least several thousand putative binding sites.

### Separation into forward and backward facing CTCF motifs

We used TFBSTools (46) to search on the 5’-3’ prime strand for forward facing CTCF motifs using the JASPAR matrix MA0139.1 and the 3’-5’ strand for motifs that are backwards facing ones. An alternative calculation using RSAT (48) to search for CTCF motifs using JASPAR matrix MA0139.1 led to similar results.

### Calculation of aggregate nucleosome profiles

Aggregate nucleosome profiles were calculated using NucTools with single-base pair resolution (43). The calculation taking into account CTCF motif directionality was done as follows: in the case, if the motif is on the plus strand the region [−1000, 1000] near CTCF also starts left to right, whereas for the minus strand the position of the region was mirrored with respect to the middle of the CTCF site.

### Stratification of TF-DNA binding affinity

In the case of experimentally determined binding sites of CTCF we stratified 33,880 sites reported by the mouse ENCODE consortium into five equally sized quintiles according to their ChIP-seq peak height reported in the original publication (9,36). In the case of computationally predicted TF sites, we have started from 111,480 sites found by scanning the mouse genome with TFBStools using JASPAR matrix MA0139.1 and split them into five equal quintiles based on their TRAP score (49) which is proportional to the binding probability of CTCF for a given site. In order to calculate the TRAP score we extended CTCF motifs by 30 nucleotides in both directions and used tRap implementation of the TRAP algorithm in R with default parameters (https://github.com/matthuska/tRap). In the calculations involving CTCF motif directionality (Figures 4-7) we first arranged predicted sites by the TRAP score into quintiles, and after that intersected them with the experimental ChIP-seq peaks of CTCF. Only motifs overlapping with sites that were experimentally detected by ChIP-seq in at least one mouse cell type were retained (including datasets from ENCODE (9), GSE27944 (50), GSE96107 (42), GSE114599 (10)), and these were further filtered to exclude CTCF sites that overlap with annotated gene promoters (which removed about 10% of CTCF sites). Promoters were defined as 1kb regions around all transcription start sites in the Genomatics Eldorado database (Genomatix GmbH). After these filtering steps we obtained the following numbers of sites in the binding strength quintiles Q1 to Q5: 3,596 (Q1); 3,782 (Q2); 6,776 (Q3); 14,776 (Q4); 16,860 (Q5).

### Phasogram calculation

The “phasograms” representing the histograms of dyad-to-dyad or start-to-start distances were calculated with the NucTools script nucleosome_repeat_length.pl. When paired-end MNase-seq was used, dyad-to-dyad distances were calculated using the center of each read as described previously (43). When chemical mapping data was used, this procedure was modified to use the start-to-start distances instead, because in the chemical mapping method the DNA cuts happen at the dyad locations, so the DNA fragments span from dyad to dyad.

### Selection of the location of the region near CTCF for NRL calculations

We noticed that NRL near CTCF depends critically on the distance of the region of NRL calculation to the binding site summit (Figure S1). While the phasograms for regions [100, 2000] and [250, 1000] near the summits of the experimental CTCF sites, which are both excluding the CTCF site, are quite similar to each other, a region that includes the peak summit [−500, 500] is characterised by a very different phasogram. However, the latter phasogram is an artefact of the effect of the interference of two “waves” of distances between nucleosomes: one wave corresponds to the distances between nucleosomes located on the same side from CTCF, and the second wave corresponds to distances between nucleosomes located on different sides from CTCF. The superposition of these two waves results in the appearance of additional peaks (Figure S1A). A linear fit through all the peaks given by the interference of these two waves gives NRL=155 bp, but this value does not reflect the real prevalent distance between nucleosomes (Figure S1B). We thus selected the region [100, 2000] for the following calculations. Below, all NRLs refer to regions [100, 2000] near the summits of TF binding sites, unless specified otherwise. We would like to note that the effect explained above means that some of the previous calculations reporting NRL near CTCF may need to be re-evaluated, because the summit of CTCF site needs to be always excluded from the genomic region for robust NRL calculations; otherwise the apparent NRL is unrealistically small. We checked that this artefact at least does not affect NRL calculations near TSS (Figure S1C), but some other previous publications may be affected. Once the region location with respect to the CTCF site is fixed, the phasograms are not significantly affected by the choice of the nucleosome positioning dataset (Figure S1D). In the following calculations in ESCs we used the high-coverage MNase-seq and chemical mapping datasets from (41).

### Automated NRL determination from phasograms

Studying many phasograms proved cumbersome when manually picking the points in a non-automated way. To circumvent this problem, an interactive applet called *NRLcalc* was developed based on the Shiny R framework (http://shiny.rstudio.com) to allow one to interactively annotate each phasogram such that the NRL could be calculated conveniently. The app allows one to select a smoothing window size to minimise noise in the phasograms. A smoothing window of 20 bp was used in our calculations. The app also provides the *Next* and *Back* button to allow the user to go through many phasograms, as well as intuitive user interface to load and save data.

## Results

### Setup of NRL calculations

Let us base our NRL calculations on the “phasogram” algorithm introduced previously (25,36). The idea of this method is to consider all mapped nucleosome reads within the genomic region of interest and calculate the distribution of the frequencies of distances between nucleosome dyads. This distribution typically shows peaks corresponding to the prevalent distance between two nearest neighbour nucleosomes followed by the distances between next neighbours. The slope of the line resulting from the linear fit of the positions of the peaks then gives the NRL (Figure 1B). To perform bulk calculations of NRLs for many genomic subsets of interest we developed software *NRLcalc*, which loads the phasograms computed in *NucTools* (43) and performs linear fitting to calculate the NRL (see Methods).

Each TF is characterised by a unique NRL distribution near its binding sites. For example, we used a recently reported chemical nucleosome mapping dataset (41) to calculate NRLs in the region of up to 2000bp from the centre of the binding site excluding the central 100 bp (hereafter referred to as region [100, 2000]) for 18 stemness-related TFs whose binding has been experimentally determined in ESCs using ChIP-seq (Figure 1C). This analysis revealed that the proximity to CTCF binding sites unanimously reduced the NRL near these sites. When we filtered out TF binding sites that overlap with CTCF binding sites in ESCs, the NRLs for each individual TF increased (Figure 1C). On the other hand, TF binding sites that overlap with CTCF had significantly smaller NRLs (Figure S2).

### The strength of CTCF binding correlates with NRL decrease in the adjacent region

To dig deeper into this newfound relationship between CTCF and local chromatin conformation, we hypothesised that CTCF binding strength would have an effect that was proportional to the decrease in NRL. To investigate this, we split CTCF sites into 5 binding strength quintiles of increasing binding strength. Two metrics were used as a means of quantifying CTCF binding strength: i) Experimentally determined CTCF binding sites in ESCs were split into 5 quintiles based on the height of the ChIP-seq peaks reported by the ENCODE consortium (9). ii) Theoretically predicted binding sites defined by scanning the mouse genome using TFBStools (46) with the 19-bp CTCF motif (JASPAR MA0139.1) (45) were split into 5 quintiles based on their calculated TRAP score that is proportional to the probability of CTCF binding to a given site (49) (see Methods). In each case, the calculation of the NRL was performed in the region [100, 2000] near CTCF binding sites using MNase-seq data (41). These calculations revealed a smooth decrease of NRL as the strength of CTCF binding increased in the case of both used metrics (Figure 2B). In addition, we used the chemical nucleosome mapping dataset (41) to compare the CTCF quintiles in terms of the distribution of nucleosome dyad-to-dyad distances, which also revealed that stronger CTCF binding is associated with smaller NRLs (Figure S3). Thus, the effect of CTCF-dependent NRL decrease is a general, dataset-independent effect. Note that chemical mapping-based NRLs should not be directly compared with MNase-seq ones due to the inherent peculiarities of the chemical mapping experiment that we noticed previously (43); below we will use only MNase-seq and ChIP-seq datasets.

**Figure 2.**
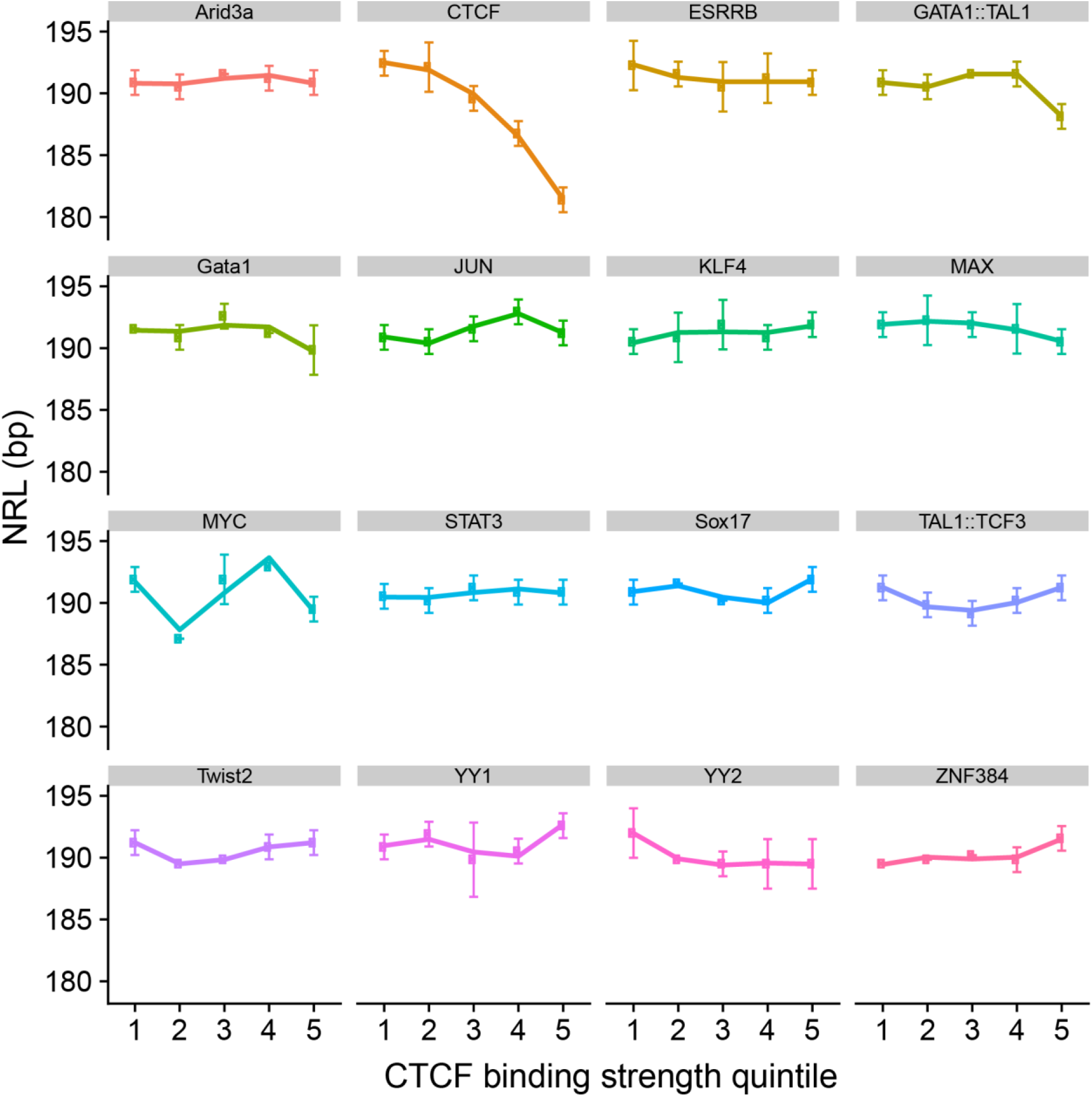
Proteins other than CTCF do not show the relationship between DNA-binding strength and NRL near their binding sites. 16 representative TFs related to stem cells are shown (similar calculations were performed for 497 TFs listed in JASPAR2018). TF bindings sites used in this analysis were predicted computationally by scanning the mouse genome using TFBStools with the 80% motif similarity cut off and then stratified into five binding strength quintiles based on the TRAP score (see methods).

Using the same procedure we have also calculated NRL in the region [100, 2000] from the TF motif as a function of the predicted TF binding strength of 497 TFs which have position weight matrices in JASPAR2018 (45). This analysis revealed that for proteins other that CTCF NRL did not reveal a smooth function of their binding strength (see Figure 2 for examples of TFs relevant to stem cells). Thus, CTCF is a unique protein whose DNA binding strength is anticorrelated to the NRL value.

### The strength of CTCF-DNA binding correlates with GC and CpG content

In order to understand the physical mechanisms of NRL decrease near CTCF we considered a number of genomic features and molecular factors that could potentially account for the NRL decrease near CTCF (Figure 3). Our previous observations suggested that the ability of CTCF site to retain CTCF during cell perturbations is related to the surrounding GC and CpG content (10,51). Our calculations performed here show that the strength of CTCF binding is indeed correlated with GC content around CTCF sites (Figure 3A), as well as the probability that a given site is located in a CpG island (Figure 3B). Furthermore, CTCF site location inside CpG islands was associated with a significantly decreased NRL in comparison with all CTCF sites (Figure 3D).

**Figure 3.**
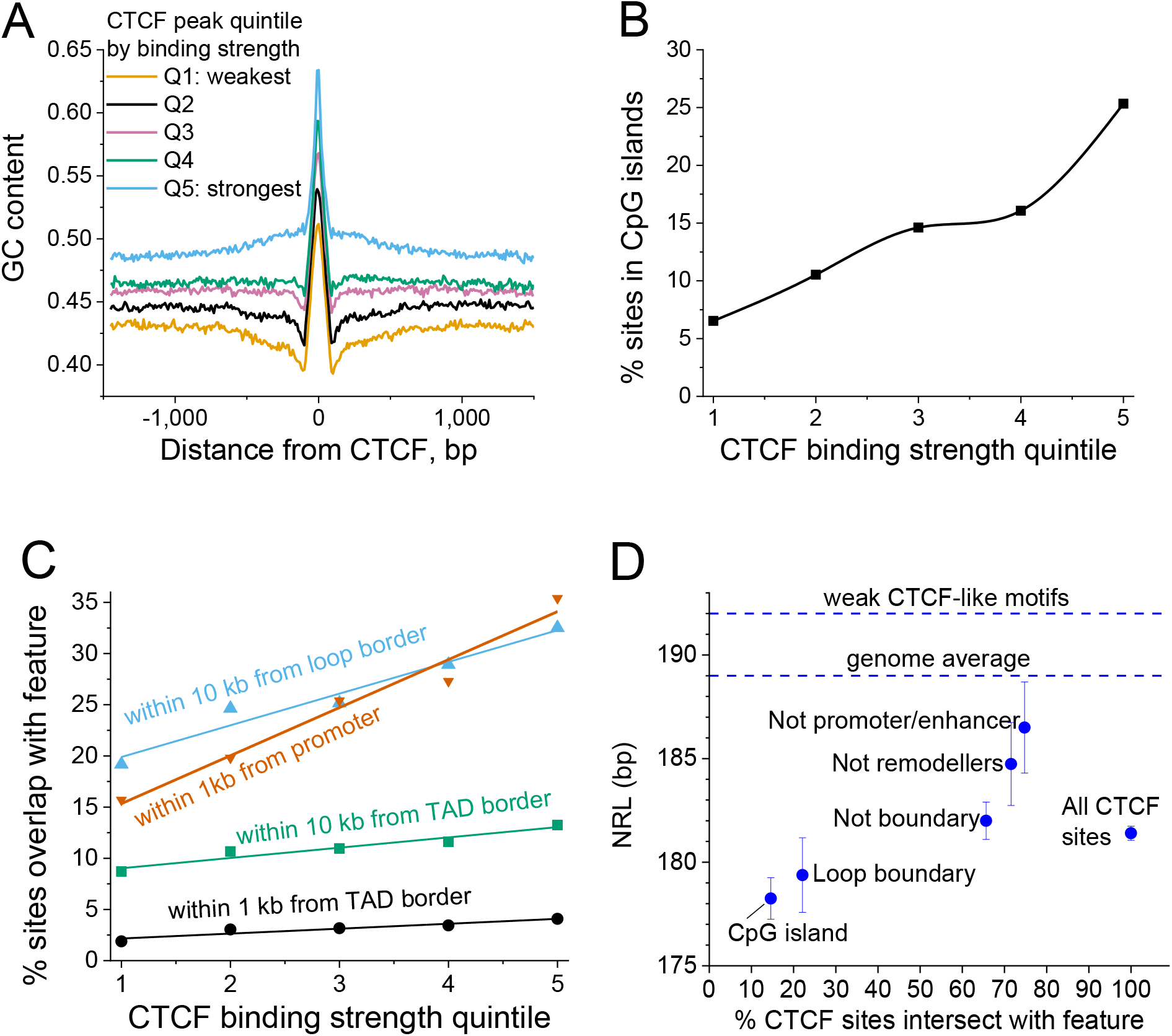
Genetic features correlating with the experimental strength of CTCF binding. A) CTCF binding sites split into quintiles based on their binding strength are characterised by increasing GC content as CTCF binding strength increases. B) The stronger CTCF binding site the higher is the probability that it is located in a CpG island. C) The stronger CTCF binds the higher the probability that it is located in a promoter or forms a boundary of TADs or enhancer-promoter loops. D) NRLs for the following subsets of CTCF sites: all sites bound in ESCs; inside chromatin loop boundary; outside of boundaries of loops and TADs; inside CpG islands; outside of chromatin remodeller peaks; outside of promoters and enhancers. The top horizontal dashed line corresponds to the weak CTCF-like motifs from Figure 2D. Vertical bars show the standard deviation.

### The strength of CTCF-DNA binding correlates with the probability of a given site to be inside cis-regulatory elements and domain boundaries

Another potential hypothesis is that the small NRL near CTCF could be because CTCF sites are in active regions (promoters, enhancers, etc.) which have a smaller NRL in comparison with genome-average based on previous studies (25,26). Our analysis performed here demonstrated that there is a positive correlation between the strength of CTCF binding and the probability that it is inside a promoter region (Figure 3C). We also used recently published coordinates of topologically associated domains (TADs) and promoter-enhancer loops in ESCs (42) and showed that there is a correlation between the strength of CTCF binding and the probability that it forms a boundary of TADs and even higher correlation for the boundaries of loops (Figure 3C). Furthermore, NRL near CTCF sites was smaller if these sites were inside borders of loops or TADs, while the NRL value went up if all known regulatory regions were excluded (Figure 3D).

### Remodeller-specific effects on NRL near CTCF

Active nucleosome positioning is determined by chromatin remodellers, but the rules of action of individual remodellers are not well defined. In order to clarify remodeller effects on NRL decrease near CTCF we processed all available remodeller ChIP-seq datasets in ESCs and plotted the percentage of CTCF sites overlapping with remodeller ChIP-seq peaks (Figure 4A). This analysis showed that the stronger CTCF binds the higher the probability that a given CTCF binding site overlaps with remodellers. Particularly large percentage of CTCF sites overlaps with peaks of remodellers Chd4, EP400, Chd8 and BRG1. Next we set to derive systematic rules of remodeller effects on NRL near CTCF (Figure 4B). By comparing NRLs near CTCF sites overlapping and non-overlapping with each remodeller, we learned that Brg1 has no detectable effect (based on two independent Brg1 datasets), and Snf2h having the strongest effect. The effect of other remodellers is increasing in the order BRG1 ≤ Chd4 < Chd6 < Chd1 ≤ Chd2 ≤ EP400 ≤ Chd8 < Snf2h (Figure 4B).

**Figure 4.**
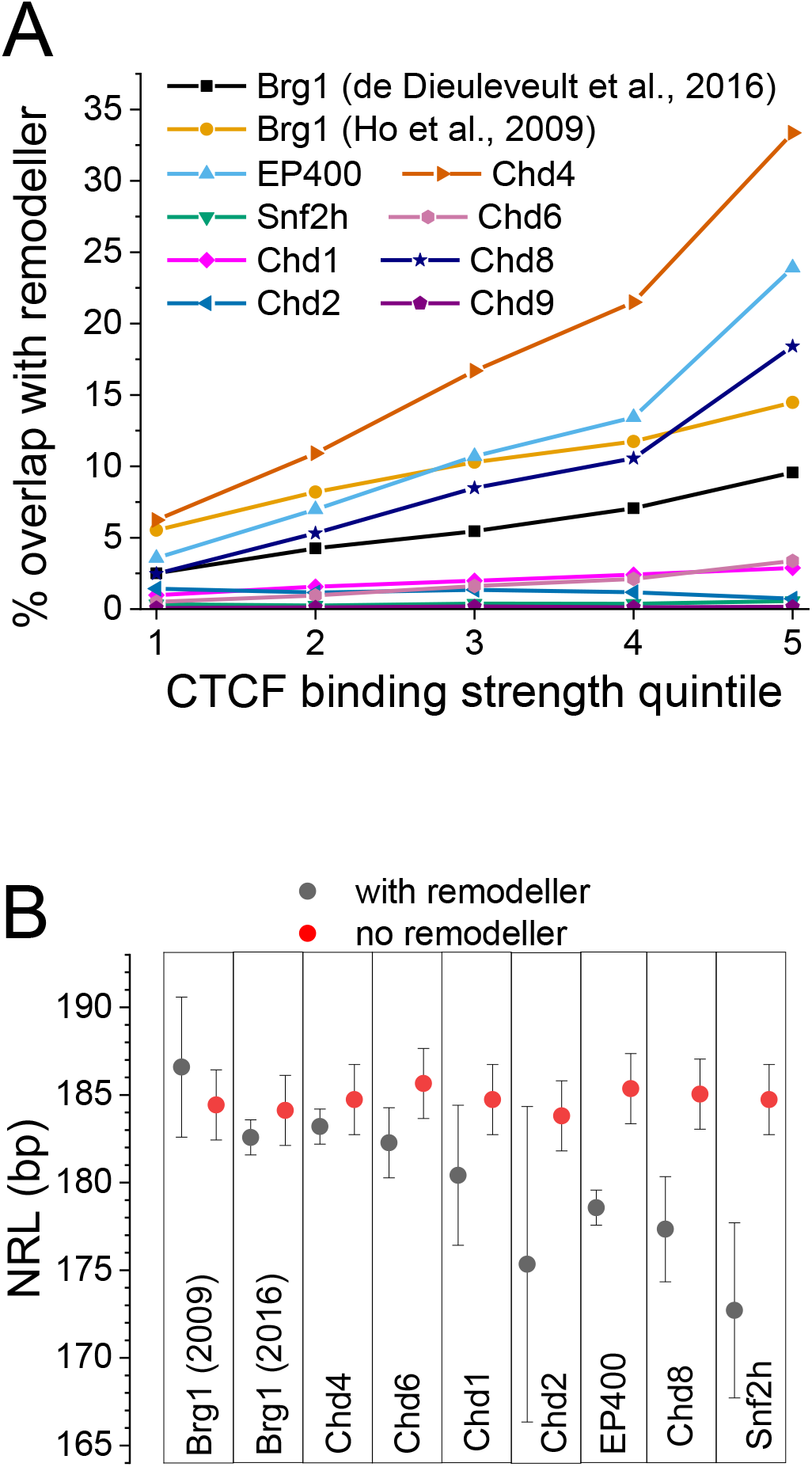
Effects of different genetic features and trans-acting factors on the value of NRL near CTCF. A) The stronger CTCF binds the higher is the probability that it is co-enriched with different chromatin remodellers indicated on the figure. The enrichment was defined as the ratio of CTCF sites overlapping with ChIP-seq peaks of a given remodeller to the total number of CTCF sites in a given quintile. B) NRLs calculated for CTCF sites that overlap (black) and do not over (red) with ChIP-seq peaks of eight chromatin remodellers experimentally mapped in ESCs. Remodeller names are indicated on the figure. Two Brg1 datasets are denoted as 2009 (73) and 2016 (34).

### CTCF motif directionality introduces asymmetry in adjacent nucleosome distribution

All our calculations above were performed without considering the directionality of the CTCF motif. For example, Figure 1A shows a symmetric pattern of nucleosome occupancy around CTCF, which arises due to averaging of different patterns around CTCF motifs in the direction of the plus and minus strand. Now let us always orient the CTCF motif in the same way, left to right (5’ to 3’), and refer to positions in 5’ direction from the CTCF motif as “upstream” and 3’ direction as “downstream”. Using this setup, we calculated aggregate profiles of nucleosome around CTCF by aligning all regions in 5’ to 3’ direction of the CTCF motif defined by the JASPAR matrix (MA0139.1). In these calculations we considered only CTCF motifs located in ChIP-seq defined peaks in at least one mouse cell type. Furthermore, we excluded CTCF sites that are located inside annotated promoters (see Methods).

Figure 5A shows the aggregate profiles of MNAse-seq nucleosome occupancy (41) around CTCF in ESCs taking into account the motif directionality. Here, the wave-like pattern of the nucleosome occupancy around CTCF sites reveals strong asymmetry. Counterintuitively, the weaker CTCF binding the stronger is the asymmetry. Such an asymmetry is similar to what is usually observed near promoters, except that we have excluded from this calculation CTCF sites that overlap with promoters. We have also confirmed this effect using MNase-assisted H3 ChIP Seq dataset (Figure S4) and plotted the occupancy of RNA Pol II around CTCF (Figure 5B). Pol II occupancy shows CTCF-dependent enrichment, which increases with the increase of CTCF binding strength. Weak CTCF sites which have the strongest asymmetry are devoid of Pol II. Thus, the asymmetry of nucleosome occupancy near CTCF is similar to the asymmetry observed for promoters, but these are not promoters and not related to Pol II-transcribed non-coding regions.

**Figure 5.**
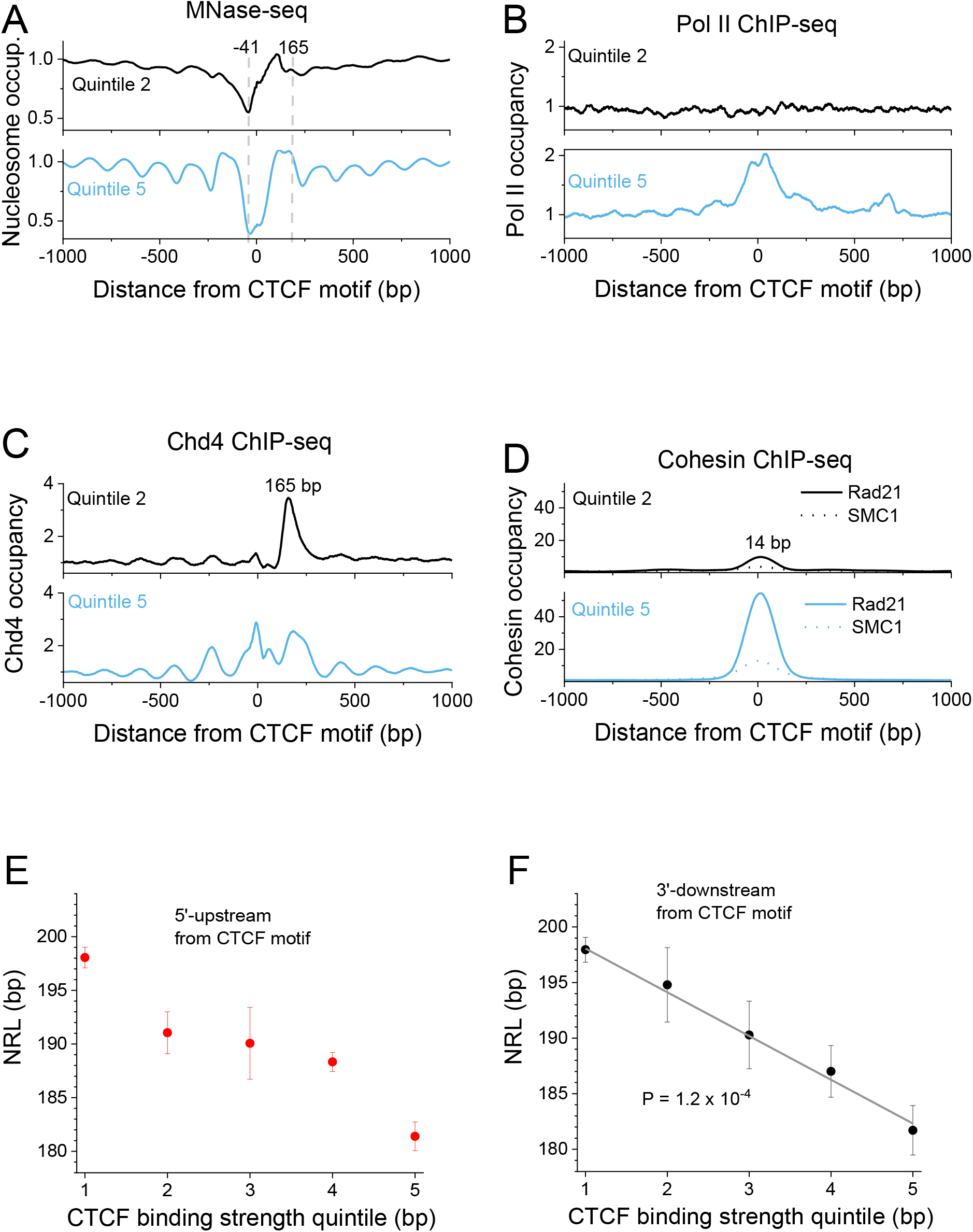
Combined effects of CTCF motif directionality and binding strength on nucleosome positioning. A) Aggregate nucleosome profiles based on MNase-seq (Voong et al) around CTCF motifs outside promoters which coincide with experimentally verified binding sites in at least one mouse cell types, taking into account the DNA strand directionality. The strong peak at 105 bp from the centre of CTCF motif appears for all CTCF quintiles. On the other hand, the nucleosome peak at position 165 is sensitive to the strength of CTCF binding and increases as the strength of CTCF binding increases from weak binding at quintile 2 to strong binding at quintile 5. B) CTCF binding outside of promoters is associated with CTCF-dependent Pol II enrichment. In the weakest CTCF quintile there is no Pol II enrichment, so the promoter-like nucleosome occupancy near CTCF is not due to Pol II. C) The binding of Chd4 (and not any other experimentally profiled remodeller) shows a CTCF dependent peak at 165 bp, coinciding with the nucleosome occupancy peak. D) The binding of cohesion subunits does not exhibit large asymmetry. The peak of Rad21 is shifted 14 bp from the centre of CTCF motif while the peak of SMC1 coincides with the centre of CTCF motif. E and F) NRL as a function of CTCF binding strength quintile corrected for the CTCF motif directionality. E) NRL calculated in the region [−2000, 100] in 5’ direction (“upstream”) of the centre of CTCF motif. F) NRL calculated in the region [100, 2000] in 3’ direction (“downstream”) of the centre of CTCF motif. In the latter case NRL dependence of CTCF binding strength can be fitted as a straight line (t-test *P* = 1.2 × 10^−4^).

The most striking feature of the asymmetric nucleosome profiles near CTCF is that the deepest point of the nucleosome-depleted region is shifted about 41 bp “upstream” in 5’ direction from the centre of the CTCF motif. This is different from what is usually assumed based on symmetric profiles such as in Figure 1A. Interestingly, the first strong nucleosome peak at 105 bp “downstream” in 3’ direction from CTCF appears similarly for all CTCF site quintiles, whereas the next peak at 165 bp “downstream” in 3’ direction from CTCF is extremely sensitive to the CTCF binding strength. There are also several other nucleosome occupancy peaks that display strong sensitivity to the CTCF binding strength. The appearance of these CTCF-dependent peaks of nucleosome occupancy as CTCF binding strength increases is causing the effect of CTCF binding strength on NRL that we observed earlier.

### The CTCF-dependent peak of nucleosome occupancy 3’-downstream of CTCF can be attributed to Chd4

In order to determine the structural origin of the peak at 165 bp from the CTCF motif we calculated aggregate profiles of all chromatin remodellers profiled using ChIP-seq in ESCs (Figure S5). Interestingly, we see in Figure S5 that the remodellers position themselves between nucleosomes. Chd4 is the only remodeller characterised by a CTCF-dependent peak at position +165 bp (Figure 5D). The peak of Chd4 at this location is quite pronounced, which suggests that while Chd4’s effect on the NRL decrease determined in Figure 4 is minor, this remodeller plays an important role in establishing the asymmetry of nucleosome positioning.

### The value of NRL in the region 3’-downstream of the CTCF motif linearly depends on the CTCF binding strength

The effect of CTCF motif directionality introduces a significant correction to the NRL dependence on the CTCF binding strength that we found above (Figure 5E and F). When performing NRL calculations separately for the region [100, 2000] 3’-downstream and region [−2000, −100] 5’-upstream from the centre of the CTCF motif, we noticed that the most regular behaviour is observed 3’-downstream where the effect can be described by a linear dependence (Figure 5F). We also checked whether the appearance of the nucleosome occupancy peak 165 bp downstream of CTCF is the main determinant of the NRL decrease. The recalculation of the NRL in the interval [300, 2000] 3’-downstream from CTCF showed that while the NRL decrease is less steep, it still follows the same trend (Figure S6).

### The asymmetric nucleosome depletion 5’-upstream of CTCF/CTCFL motifs is encoded in DNA repeats and may be linked to their transcription

Next we calculated the average nucleotide distribution around CTCF sites used above taking into account the orientation of CTCF motifs. This revealed an unexpected nucleotide pattern in the extended region near CTCF (Figure 6). The nucleosome depletion in the region around −41 bp upstream of CTCF is associated with a decrease of GC content. This is consistent with previous observations that high AT-content and in particular poly(dA:dT)-tracts have strong nucleosome-excluding properties (52). It is worth noting that the CTCF motif used in our calculations is just 19 bp, but the length of the highly structured area near CTCF is more than 200 bp. This means that the CTCF motif is frequently encountered as part of a much larger DNA sequence organisation, some type of sequence repeats that are primarily responsible for the establishment of the asymmetric boundaries around CTCF. Indeed, 50% of the CTCF motifs used in our calculations in Figures 5 and 6 overlapped with repeats defined by the UCSC Genome Browser repeat masker. Furthermore, the percentage of repeats given by the repeat masker shows a similar very structured profile with an extended region (>200 bp) near CTCF strongly enriched with repeats.

**Figure 6.**
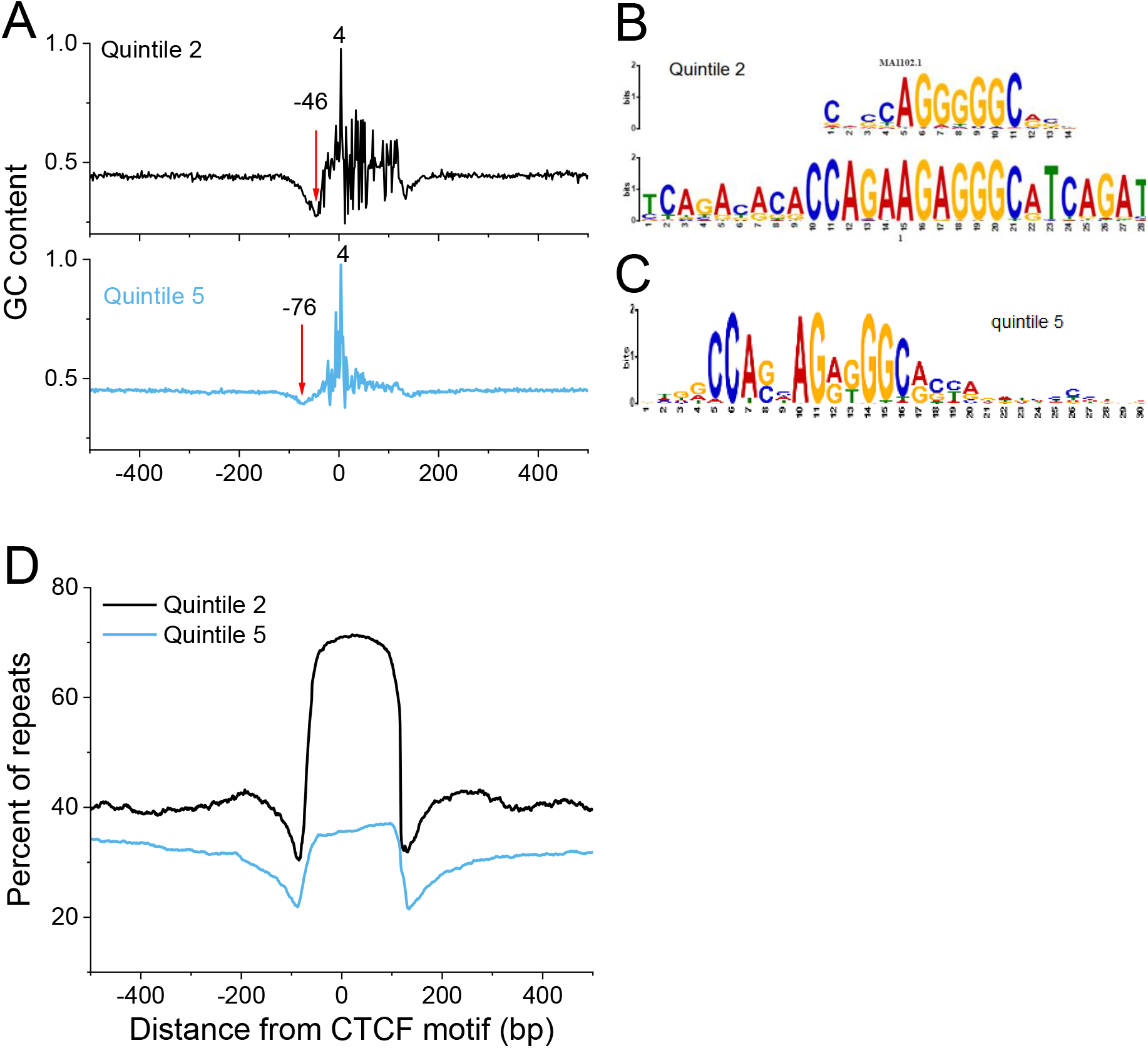
Effects of the nucleotide content around CTCF sites. A) Average GC content around CTCF motifs for CTCF binding strength quintiles 2 and 5. B) The sequence of the consensus motif in quintile 2 with the smallest P-value. The best TF match for the quintile 2 consensus motif is CTCFL (Boris) (JASPAR MA1102.1). C) The sequence of the consensus motif in quintile 5. The quintile 5 consensus sequence contains the classical CTCF motif (JASPAR MA0139.1). D) The percentage of repeats determined by the USCS Genome Browser’s Repeat Masker as a function of the distance from the middle of CTCF motifs.

We have also checked whether the nucleosome depletion 5’-upstream of CTCF is related to transposon transcription. Using coordinates of ChIP-seq peaks of RNA Pol III determined previously in ESCs (53), we found that 33% of co-localisations of TFIIIC and Pol III and 17% of co-localisations of SINE repeats and Pol III overlapped with our CTCF motifs. Thus, not only the DNA repeats are responsible for the AT-rich region 5’-upstream of CTCF, but also their transcription may be linked to the asymmetric nucleosome depletion pattern.

Another interesting finding shown in Figure 6B and C is that when we subjected each binding strength quintile to a separate de novo motif discovery, the strongest quintile 5 was associated with the classical CTCF motif (JASPAR MA0139.1), whereas a weak quintile 2 was associated with CTCFL (BORIS) defined by the JASPAR matrix MA1102.1.

### Nucleosome-depleted boundaries 5’-upstream of CTCF motif are preserved even if binding CTCF is lost during cell differentiation

Next we compared nucleosome positioning around CTCF motifs upon differentiation of ESCs to neural progenitor cells (NPSs) and mouse embryonic fibroblasts (MEFs) using MNase-seq data from (36) and CTCF ChIP-seq data from (9,42) (Figure 7A). Notably, stronger CTCF binding to DNA increases the probability that a given site will remain bound upon differentiation. This suggests that the sequence-dependent strength of CTCF binding can act as the “CTCF code”, determining which CTCF sites retain and which are lost upon differentiation (and thus how the 3D structure of the genome will change). Our further analysis revealed that common CTCF sites that are present in all three states are characterised by quite minor asymmetry of nucleosome organisation (Figure 7B). On the other hand, CTCF sites that are lost upon ESC differentiation to NPCs and MEFs have more profound asymmetry of the nucleosome pattern around them (Figure 7C and D). Upon differentiation both in NPCs and MEFs, the array of nucleosome 3’-downstream of the CTCF motif is shifted to cover the CTCF site. Interestingly, the nucleosome-depleted region 5’-upstream of CTCF still remains open upon differentiation. The latter effect was also confirmed for the case of CTCF sites that are not bound by CTCF in ESCs and become bound in MEFs (Figure S7).

**Figure 7.**
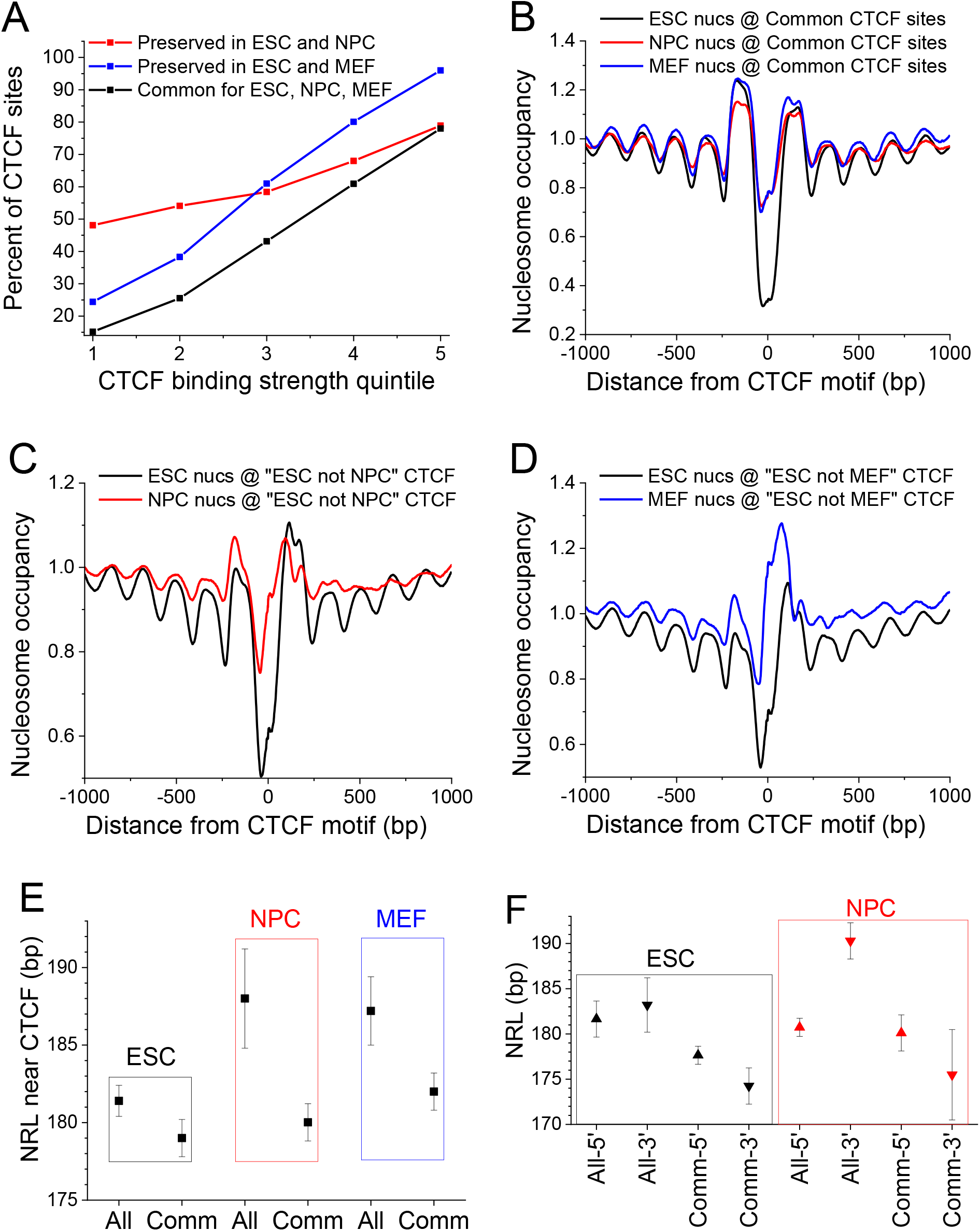
Effects of asymmetric CTCF-dependent boundaries in stem cell differentiation. A) The fraction of CTCF sites preserved upon differentiation of ESCs to NPCs and MEFs as a function of CTCF binding strength. CTCF sites preserves in all these three cell types are termed “common”. B) Nucleosome occupancy in ESCs (black), NPCs (red) and MEFs (blue) around CTCF sites common between ESC, NPC and MEF, calculated taking into account CTCF motif directionality. C) Nucleosome occupancy around “ESC not MEF” sites that are present in ESCs (black line) but lost in MEFs (red line) taking into account CTCF motif directionality. D) Nucleosome occupancy around “ESC not NPC” sites that are present in ESCs (black line) but lost in NPCs (red line) taking into account CTCF motif directionality. Note that in differentiated cells a nucleosome is being positioned to cover the “lost” CTCF sites, but nucleosome depletion on the left of CTCF is still preserved. E) NRLs in region [100, 2000] from CTCF’s experimental binding site summit calculated without taking into account the motif directionality. Upon differentiation average NRL near CTCF increases (denoted “All”), but common CTCF sites keep the smallest NRL (denoted “Comm”). F) NRLs in region [100, 2000] from CTCF’s binding motifs overlapping with experimentally confirmed CTCF binding sites, calculated separately 5’-upstream and 3’-downstream of CTCF motifs. The main NRL change during differentiation is in the region 3’-downstream of CTCF motifs.

### Common CTCF sites preserve local nucleosome organisation during ESC differentiation

Then, we set to determine the functional consequences of the NRL decrease near CTCF. NRL near bound CTCF on average increases as the cell differentiates from ESCs to NPCs or MEFs (Figure 7E and S8). However, common CTCF sites resist this NRL change, suggesting that CTCF retention at common sites upon differentiation preserves both 3D structure and nucleosome patterns at these loci. As we have established previously (Figure 5F), the effect of the active CTCF-dependent NRL decrease is mostly pronounced 3’-downstream of CTCF motifs. The NRL increase near CTCF upon cell differentiation is also mostly in the 3’-downstream region (Figure 7F).

## Discussion

We developed a new *NRLcalc* methodology to investigate nucleosome rearrangement and NRL changes near TF binding motifs distinguished by their orientation and binding strength, and the application of this method to CTCF revealed a number of new effects (Figure 8):

**Figure 8.**
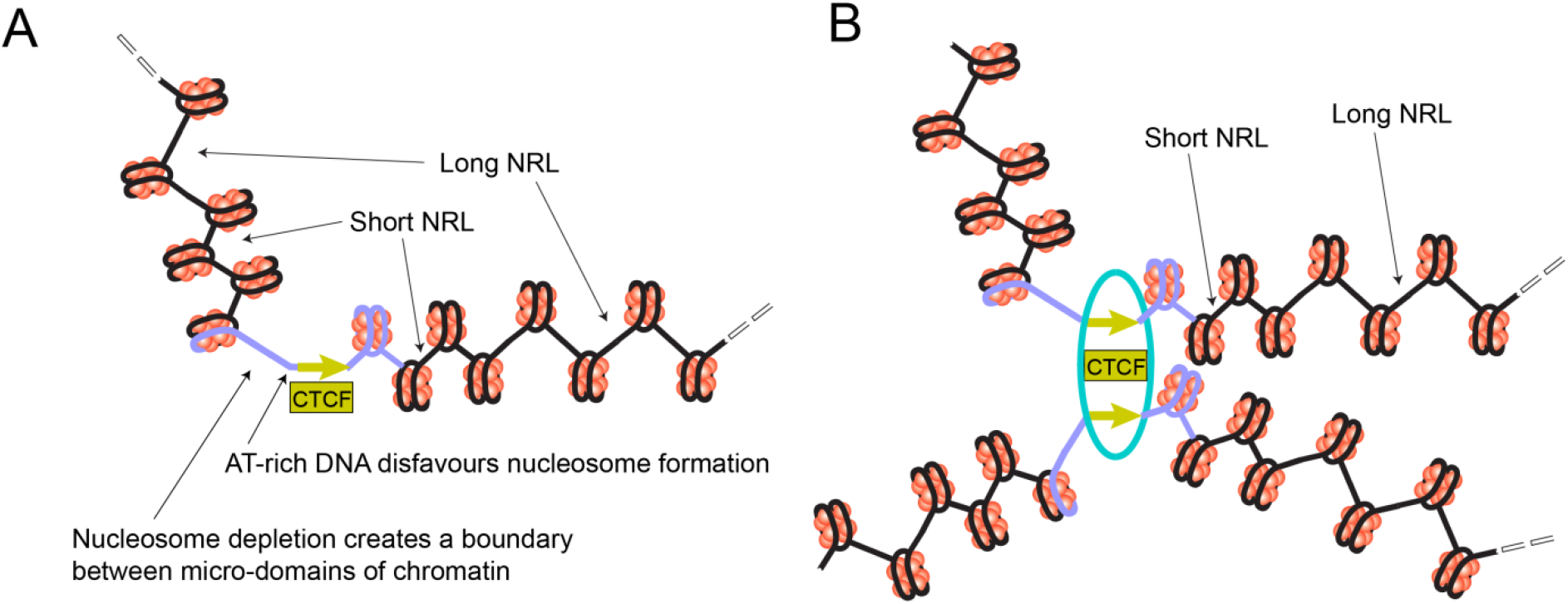
Schematic illustration of the effect of CTCF binding strength and motif orientation on the nucleosome arrangement in a single genomic region (A) and at the base of a loop (B). An extended DNA region including CTCF motif is enriched with repetitive sequences that define the mechanical properties of this region as a chromatin boundary (shown in violet colour) – see figures 5A, 6A, 6D and S4. The region 5’-upstream of CTCF motif contains AT-rich sequences that disfavour nucleosome formation and may account for DNA bending in the complex with CTCF. Such regions can be due to DNA repeats such as SINEs, some of which are transcribed by Pol III that interact with CTCF. In analogy to the coding gene transcription the region 5’-upstream of the CTCF motif is depleted of the “−1” nucleosome. In the region 3’-downstream of CTCF motif chromatin remodellers including Chd4 and Snf2h determine the regularity of the nucleosome array. The nucleosomes located close to CTCF are separated by shorter linkers and nucleosomes further away from CTCF are separated by longer linkers, reaching the genome-average linker length at distances where CTCF effects disappear (corresponding to NRL change from ∼180 bp near strong CTCF sites to ∼190 bp genome-average, see Figure 3D). The cohesin ring is represented by the cyan ellipse.

Firstly, we found that contrary to previous assumptions, the nucleosome arrangement near CTCF motifs is asymmetric and to a large degree hard-wired in the sequence of the DNA region >200 bp long including the CTCF motif (Figure 5A and 6A). The asymmetry in this case is not just a consequence of heterogeneity of nucleosome distributions around subsets of sites (54), but is a generic feature across all CTCF sites. The nucleosome-depleted region, which was previously believed to coincide with the CTCF binding site (37,38), is actually shifted 5’-upstream of CTCF motif (Figure 5). This nucleosome depletion is associated with AT-rich DNA sequence repeats which may disfavour nucleosome formation (52) and introduce bending of the double helix near CTCF (55,56). We showed here that these regions may be linked to transcription of transposons such as Pol III-dependent SINE repeats. Several publications suggested important roles of transposons in the evolution of CTCF sites (57–61), and also it is known that mouse SINE B2 repeats can act as insulators (domain boundaries) *per se* (62). In addition, our data suggests that CTCF may play active role in transposon functioning as transcribed units separating nucleosome arrays. Interestingly, previous publications reported that TFIIIC binds to RNA Pol III at tRNA genes and acts as a barrier against the spreading of heterochromatin (63) – this barrier function can be now re-interpreted in light of our results on the association of CTCF with Pol III as well as Pol II outside of gene promoters (Figure 5B).

We also showed that the asymmetry of the nucleosome signatures depends on the DNA-defined strength of CTCF binding and may be in addition determined by the CTCF/CTCFL competition, because “weak” CTCF binding sites are enriched with the CTCFL recognition motif (Figure 6). CTCFL, also known as BORIS, has been previously proposed to interfere with CTCF binding (64), and our results further substantiate its role in the “CTCF code” (42) that defines differential CTCF/CTCFL binding.

Secondly, we found that the NRL decrease near CTCF is correlated with CTCF-DNA binding affinity (Figure 1D and 5F). This result goes significantly beyond previous observations that the CTCF binding strength is related to a more regular nucleosome ordering near its binding site (43,65) and may have direct functional implications. Strikingly, the variation of NRL as a function of CTCF binding affinity can be as large as ∼20 bp (the difference between NRL near the weakest CTCF-like motifs and the strongest CTCF-bound sites). None of other DNA-binding proteins showed such behaviour (Figure 2). This uniqueness of CTCF can be explained by the large variability of its binding affinity through different combinations of its 11 zinc fingers that allows creating a “CTCF code” (56,64). The effect of the NRL dependence on CTCF binding strength is most profound 3’-downstream of CTCF motifs, where it can be approximated by a linear function (Figure 5F). This strong nucleosome patterning downstream but not upstream of CTCF is comparable to that of transcription start sites (TSSs) of protein-coding genes. In analogy, this effect could provide an additional argument that this may be linked to the transcription of non-coding repeats enclosing CTCF including Pol III-dependent SINEs.

Thus, our data suggests that the NRL decrease near CTCF is a result of an active, remodeller-dependent process. Therefore, we analysed the contributions to NRL decrease caused by each of 8 chromatin remodellers that have been profiled in ESCs (Fig 4B). We found that that Snf2h has a major role in this phenomenon, consistent with previous studies of Snf2H knockout in HeLa cells (66) and ESCs (39). In accord with the latter study, we observed that BRG1 has no detectable effect on NRL near CTCF, although it may be still involved in nucleosome positioning near TAD boundaries (67). Our investigation also identified Chd8 and EP400 as two major players on nucleosome arrangement near CTCF (Figure 4B, S5G). These findings are consistent with the previous investigations that showed that Chd8 physically interacts with CTCF and knockdown of Chd8 abolishes the insulator activity of CTCF sites required for *IGF2* imprinting (68). One can hypothesise that this kind of insulator activity of CTCF is related to the boundary created by the nucleosome-free region 5’-upstream of the CTCF motif reported here, which may physically prevent the spreading of DNA methylation and other epigenetic modifications. According to our analysis, the main chromatin remodeller responsible for the asymmetry of the nucleosome array near CTCF is Chd4. We show that Chd4 is the sole remodeller responsible for the CTCF-dependent nucleosome occupancy peak 3’-downstream of CTCF (Figure 5C). Interestingly, recent studies indicated that Chd4 is increasing the nucleosome density at regulatory regions (69).

Finally, we investigated the effects of CTCF motif directionality and binding strength on nucleosome rearrangement during cell differentiation. Our calculations showed that the binding affinity is a good predictor for a given CTCF site being preserved upon cell differentiation (Figure 7A). This may be used as a foundation for the “CTCF code” determining its differential binding as the cell progresses along the Waddington-type pathways. A specific subclass of common CTCF sites preserved upon cell differentiation tends to keep a small NRL, while the average NRL near all CTCF sites increases due to the active nucleosome repositioning 3’-downstream of CTCF motifs (Figure 7). A previous study reported a related distinction of common versus non-common CTCF sites based on the distance between the two nucleosomes downstream and upstream of CTCF (70). The preservation of NRL for common CTCF sites may give rise to a new effect where differential CTCF binding defines extended regions which do not change (or change minimally) their nucleosome positioning. Unexpectedly, the nucleosome-depleted region 5’-upstream of the CTCF motif remains even after CTCF depletion from a given site during differentiation. These nucleosome-depleted regions can have important functional roles, including the preservation of chromatin states while CTCF-dependent loops are dynamic and frequently break and reform throughout the cell cycle (71). For example, if the spreading of some chemical modifications of DNA or histones along the genomic coordinate requires enzymes cooperatively binding to the adjacent nucleosomes, then the consistent lack of a nucleosome at a given location can stop the propagation of the “epigenetic wave”.

Our finding of the asymmetry of CTCF-dependent chromatin boundaries at the scale of several nucleosomes may also provide the missing mechanistic explanation for the asymmetry of chromatin boundaries at the scale of hundreds to thousands of nucleosomes in so called “stripe” chromatin domains reported recently (72). In general, the asymmetric nucleosome organisation near CTCF reported here can be particularly interesting in light of the ongoing debate on the functional roles of chromatin boundaries in gene regulation.

## Data availability

Our software is available at https://github.com/chrisclarkson/NRLcalc

## Supplementary Data

Supplementary materials include Figures S1-S7 and Table ST1.

## Acknowledgements

We thank Boyan Bonev and Giacomo Cavalli for providing the coordinates of chromatin loops and TADs, Feng Cui and Noam Kaplan for helpful discussions, Stuart Newman for the computer cluster support and Yevhen Vainshtein for the NucTools support.

## Author contributions

Study design: CTC, VBZ, TVB; Calculations: CTC, EAD, RS, HM, VBZ, TVB; Manuscript draft: VBZ, TVB.

## References

1. Teif, V.B. and Clarkson, C.T. (2019) In Ranganathan, S., Gribskov, M., Nakai, K. and Schönbach, C. (eds.), Encyclopedia of Bioinformatics and Computational Biology. Academic Press, Oxford, pp. 308–317.

2. Baldi, S. (2019) Nucleosome positioning and spacing: from genome-wide maps to single arrays. Essays In Biochemistry, EBC20180058.

3. Lai, W.K.M. and Pugh, B.F. (2017) Understanding nucleosome dynamics and their links to gene expression and DNA replication. Nat Rev Mol Cell Biol, 18, 548–562.

4. Maeshima, K., Ide, S. and Babokhov, M. (2019) Dynamic chromatin organization without the 30-nm fiber. Current Opinion in Cell Biology, 58, 95–104.

5. Merkenschlager, M. and Nora, E.P. (2016) CTCF and Cohesin in Genome Folding and Transcriptional Gene Regulation. Annu Rev Genomics Hum Genet, 17, 17–43.

6. Rao, S.S.P., Huang, S.-C., Glenn St Hilaire, B., Engreitz, J.M., Perez, E.M., Kieffer-Kwon, K.-R., Sanborn, A.L., Johnstone, S.E., Bascom, G.D., Bochkov, I.D. et al. (2017) Cohesin Loss Eliminates All Loop Domains. Cell, 171, 305–320.e324.

7. Nora, E.P., Goloborodko, A., Valton, A.L., Gibcus, J.H., Uebersohn, A., Abdennur, N., Dekker, J., Mirny, L.A. and Bruneau, B.G. (2017) Targeted Degradation of CTCF Decouples Local Insulation of Chromosome Domains from Genomic Compartmentalization. Cell, 169, 930–944.

8. Fu, Y., Sinha, M., Peterson, C.L. and Weng, Z. (2008) The insulator binding protein CTCF positions 20 nucleosomes around its binding sites across the human genome. PLoS Genetics, 4, e1000138.

9. Shen, Y., Yue, F., McCleary, D.F., Ye, Z., Edsall, L., Kuan, S., Wagner, U., Dixon, J., Lee, L., Lobanenkov, V.V. et al. (2012) A map of the cis-regulatory sequences in the mouse genome. Nature, 488, 116–120.

10. Wiehle, L., Thorn, G.J., Raddatz, G., Clarkson, C.T., Rippe, K., Lyko, F., Breiling, A. and Teif, V.B. (2019) DNA (de)methylation in embryonic stem cells controls CTCF-dependent chromatin boundaries. Genome Res, 29, 750–761.

11. Wang, H., Maurano, M.T., Qu, H., Varley, K.E., Gertz, J., Pauli, F., Lee, K., Canfield, T., Weaver, M., Sandstrom, R. et al. (2012) Widespread plasticity in CTCF occupancy linked to DNA methylation. Genome Res, 22, 1680–1688.

12. Chen, H., Tian, Y., Shu, W., Bo, X. and Wang, S. (2012) Comprehensive identification and annotation of cell type-specific and ubiquitous CTCF-binding sites in the human genome. PLoS One, 7, e41374.

13. Routh, A., Sandin, S. and Rhodes, D. (2008) Nucleosome repeat length and linker histone stoichiometry determine chromatin fiber structure. Proc Natl Acad Sci U S A, 105, 8872–8877.

14. Bass, M.V., Nikitina, T., Norouzi, D., Zhurkin, V.B. and Grigoryev, S.A. (2019) Nucleosome spacing periodically modulates nucleosome chain folding and DNA topology in circular nucleosome arrays. J Biol Chem, 294, 4233–4246.

15. Bascom, G.D., Kim, T. and Schlick, T. (2017) Kilobase Pair Chromatin Fiber Contacts Promoted by Living-System-Like DNA Linker Length Distributions and Nucleosome Depletion. J Phys Chem B, 121, 3882–3894.

16. Risca, V.I., Denny, S.K., Straight, A.F. and Greenleaf, W.J. (2017) Variable chromatin structure revealed by in situ spatially correlated DNA cleavage mapping. Nature, 541, 237–241.

17. Nikitina, T., Norouzi, D., Grigoryev, S.A. and Zhurkin, V.B. (2017) DNA topology in chromatin is defined by nucleosome spacing. Science Advances, 3, e1700957.

18. Olins, A.L. and Olins, D.E. (1974) Spheroid chromatin units (v bodies). Science, 183, 330–332.

19. Kornberg, R.D. (1974) Chromatin structure: a repeating unit of histones and DNA. Science, 184, 868–871.

20. Lohr, D., Tatchell, K. and Van Holde, K.E. (1977) On the occurrence of nucleosome phasing in chromatin. Cell, 12, 829–836.

21. Gottesfeld, J.M. and Melton, D.A. (1978) The length of nucleosome-associated DNA is the same in both transcribed and nontranscribed regions of chromatin. Nature, 273, 317–319.

22. De Ambrosis, A., Ferrari, N., Bonassi, S. and Vidali, G. (1987) Nucleosomal repeat length in active and inactive genes. FEBS Lett, 225, 120–122.

23. Weintraub, H. (1978) The nucleosome repeat length increases during erythropoiesis in the chick. Nucleic Acids Res, 5, 1179–1188.

24. van Holde, K.E. (1989) Chromatin. Springer-Verlag, New York.

25. Valouev, A., Johnson, S.M., Boyd, S.D., Smith, C.L., Fire, A.Z. and Sidow, A. (2011) Determinants of nucleosome organization in primary human cells. Nature, 474, 516–520.

26. Baldi, S., Krebs, S., Blum, H. and Becker, P.B. (2018) Genome-wide measurement of local nucleosome array regularity and spacing by nanopore sequencing. Nat Struct Mol Biol, 25, 894–901.

27. Chereji, R.V., Ramachandran, S., Bryson, T.D. and Henikoff, S. (2018) Precise genome-wide mapping of single nucleosomes and linkers in vivo. Genome Biol, 19, 19.

28. Sun, F.-L., Cuaycong, M.H. and Elgin, S.C.R. (2001) Long-Range Nucleosome Ordering Is Associated with Gene Silencing in *Drosophila melanogaster* Pericentric Heterochromatin. Molecular and Cellular Biology, 21, 2867–2879.

29. Zhang, Z., Wippo, C.J., Wal, M., Ward, E., Korber, P. and Pugh, B.F. (2011) A packing mechanism for nucleosome organization reconstituted across a eukaryotic genome. Science, 332, 977–980.

30. Hennig, B.P., Bendrin, K., Zhou, Y. and Fischer, T. (2012) Chd1 chromatin remodelers maintain nucleosome organization and repress cryptic transcription. EMBO Rep, 13, 997–1003.

31. Kubik, S., Challal, D., Bruzzone, M.J., Dreos, R., Mattarocci, S., Bucher, P., Libri, D. and Shore, D. (2019) Opposing chromatin remodelers control transcription initiation frequency and start site selection. bioRxiv, 592816.

32. Ocampo, J., Chereji, R.V., Eriksson, P.R. and Clark, D.J. (2016) The ISW1 and CHD1 ATP-dependent chromatin remodelers compete to set nucleosome spacing in vivo. Nucleic Acids Res, 44, 4625–4635.

33. Mobius, W., Osberg, B., Tsankov, A.M., Rando, O.J. and Gerland, U. (2013) Toward a unified physical model of nucleosome patterns flanking transcription start sites. Proc Natl Acad Sci U S A, 110, 5719–5724.

34. de Dieuleveult, M., Yen, K., Hmitou, I., Depaux, A., Boussouar, F., Bou Dargham, D., Jounier, S., Humbertclaude, H., Ribierre, F., Baulard, C. et al. (2016) Genome-wide nucleosome specificity and function of chromatin remodellers in ES cells. Nature, 530, 113–116.

35. Giles, K.A., Gould, C.M., Du, Q., Skvortsova, K., Song, J.Z., Maddugoda, M.P., Achinger-Kawecka, J., Stirzaker, C., Clark, S.J. and Taberlay, P.C. (2019) Integrated epigenomic analysis stratifies chromatin remodellers into distinct functional groups. Epigenetics Chromatin, 12, 12.

36. Teif, V.B., Vainshtein, Y., Caudron-Herger, M., Mallm, J.P., Marth, C., Höfer, T. and Rippe, K. (2012) Genome-wide nucleosome positioning during embryonic stem cell development. Nat Struct Mol Biol, 19, 1185–1192.

37. Teif, V.B., Beshnova, D.A., Vainshtein, Y., Marth, C., Mallm, J.P., Höfer, T. and Rippe, K. (2014) Nucleosome repositioning links DNA (de)methylation and differential CTCF binding during stem cell development. Genome Res, 24, 1285–1295.

38. Beshnova, D.A., Cherstvy, A.G., Vainshtein, Y. and Teif, V.B. (2014) Regulation of the nucleosome repeat length in vivo by the DNA sequence, protein concentrations and long-range interactions. PLoS Comput Biol, 10, e1003698.

39. Barisic, D., Stadler, M.B., Iurlaro, M. and Schübeler, D. (2019) Mammalian ISWI and SWI/SNF selectively mediate binding of distinct transcription factors. Nature.

40. Jenkinson, G., Pujadas, E., Goutsias, J. and Feinberg, A.P. (2017) Potential energy landscapes identify the information-theoretic nature of the epigenome. Nat Genet, 49, 719–729.

41. Voong, L.N., Xi, L., Sebeson, A.C., Xiong, B., Wang, J.P. and Wang, X. (2016) Insights into Nucleosome Organization in Mouse Embryonic Stem Cells through Chemical Mapping. Cell, 167, 1555–1570 e1515.

42. Bonev, B., Mendelson Cohen, N., Szabo, Q., Fritsch, L., Papadopoulos, G.L., Lubling, Y., Xu, X., Lv, X., Hugnot, J.P., Tanay, A. et al. (2017) Multiscale 3D Genome Rewiring during Mouse Neural Development. Cell, 171, 557–572 e524.

43. Vainshtein, Y., Rippe, K. and Teif, V.B. (2017) NucTools: analysis of chromatin feature occupancy profiles from high-throughput sequencing data. BMC Genomics, 18, 158.

44. Quinlan, A.R. (2014) BEDTools: The Swiss-Army Tool for Genome Feature Analysis. Curr Protoc Bioinformatics, 47, 11 12 11–34.

45. Khan, A., Fornes, O., Stigliani, A., Gheorghe, M., Castro-Mondragon, J.A., van der Lee, R., Bessy, A., Cheneby, J., Kulkarni, S.R., Tan, G. et al. (2018) JASPAR 2018: update of the open-access database of transcription factor binding profiles and its web framework. Nucleic Acids Res, 46, D1284.

46. Tan, G. and Lenhard, B. (2016) TFBSTools: an R/bioconductor package for transcription factor binding site analysis. Bioinformatics, 32, 1555–1556.

47. Lawrence, M., Huber, W., Pages, H., Aboyoun, P., Carlson, M., Gentleman, R., Morgan, M.T. and Carey, V.J. (2013) Software for computing and annotating genomic ranges. PLoS Comput Biol, 9, e1003118.

48. Castro-Mondragon, J.A., Jaeger, S., Thieffry, D., Thomas-Chollier, M. and van Helden, J. (2017) RSAT matrix-clustering: dynamic exploration and redundancy reduction of transcription factor binding motif collections. Nucleic Acids Res, 45, e119.

49. Roider, H.G., Kanhere, A., Manke, T. and Vingron, M. (2007) Predicting transcription factor affinities to DNA from a biophysical model. Bioinformatics, 23, 134–141.

50. Martin, D., Pantoja, C., Fernandez Minan, A., Valdes-Quezada, C., Molto, E., Matesanz, F., Bogdanovic, O., de la Calle-Mustienes, E., Dominguez, O., Taher, L. et al. (2011) Genome-wide CTCF distribution in vertebrates defines equivalent sites that aid the identification of disease-associated genes. Nat Struct Mol Biol, 18, 708–714.

51. Pavlaki, I., Docquier, F., Chernukhin, I., Kita, G., Gretton, S., Clarkson, C.T., Teif, V.B. and Klenova, E. (2018) Poly(ADP-ribosyl)ation associated changes in CTCF-chromatin binding and gene expression in breast cells. Biochim Biophys Acta Gene Regul Mech, 1861, 718–730.

52. Segal, E. and Widom, J. (2009) Poly(dA:dT) tracts: major determinants of nucleosome organization. Curr Opin Struct Biol, 19, 65–71.

53. Carriere, L., Graziani, S., Alibert, O., Ghavi-Helm, Y., Boussouar, F., Humbertclaude, H., Jounier, S., Aude, J.C., Keime, C., Murvai, J. et al. (2012) Genomic binding of Pol III transcription machinery and relationship with TFIIS transcription factor distribution in mouse embryonic stem cells. Nucleic Acids Res, 40, 270–283.

54. Kundaje, A., Kyriazopoulou-Panagiotopoulou, S., Libbrecht, M., Smith, C.L., Raha, D., Winters, E.E., Johnson, S.M., Snyder, M., Batzoglou, S. and Sidow, A. (2012) Ubiquitous heterogeneity and asymmetry of the chromatin environment at regulatory elements. Genome Res, 22, 1735–1747.

55. Ghirlando, R. and Felsenfeld, G. (2016) CTCF: making the right connections. Genes Dev, 30, 881–891.

56. Nichols, M.H. and Corces, V.G. (2015) A CTCF Code for 3D Genome Architecture. Cell, 162, 703–705.

57. Zhang, Y., Li, T., Preissl, S., Grinstein, J., Farah, E., Destici, E., Lee, A.Y., Chee, S., Qiu, Y., Ma, K. et al. (2019) 3D Chromatin Architecture Remodeling during Human Cardiomyocyte Differentiation Reveals A Role Of HERV-H In Demarcating Chromatin Domains. bioRxiv, 485961.

58. Choudhary, M.N., Friedman, R.Z., Wang, J.T., Jang, H.S., Zhuo, X. and Wang, T. (2018) Co-opted transposons help perpetuate conserved higher-order chromosomal structures. bioRxiv, 485342.

59. Kentepozidou, E., Aitken, S.J., Feig, C., Stefflova, K., Ibarra-Soria, X., Odom, D.T., Roller, M. and Flicek, P. (2019) Clustered CTCF binding is an evolutionary mechanism to maintain topologically associating domains. bioRxiv, 668855.

60. Schmidt, D., Schwalie, P.C., Wilson, M.D., Ballester, B., Goncalves, A., Kutter, C., Brown, G.D., Marshall, A., Flicek, P. and Odom, D.T. (2012) Waves of retrotransposon expansion remodel genome organization and CTCF binding in multiple mammalian lineages. Cell, 148, 335–348.

61. Bourque, G., Leong, B., Vega, V.B., Chen, X., Lee, Y.L., Srinivasan, K.G., Chew, J.L., Ruan, Y., Wei, C.L., Ng, H.H. et al. (2008) Evolution of the mammalian transcription factor binding repertoire via transposable elements. Genome Res, 18, 1752–1762.

62. Lunyak, V.V., Prefontaine, G.G., Nunez, E., Cramer, T., Ju, B.G., Ohgi, K.A., Hutt, K., Roy, R., Garcia-Diaz, A., Zhu, X. et al. (2007) Developmentally regulated activation of a SINE B2 repeat as a domain boundary in organogenesis. Science, 317, 248–251.

63. Simms, T.A., Dugas, S.L., Gremillion, J.C., Ibos, M.E., Dandurand, M.N., Toliver, T.T., Edwards, D.J. and Donze, D. (2008) TFIIIC binding sites function as both heterochromatin barriers and chromatin insulators in Saccharomyces cerevisiae. Eukaryotic cell, 7, 2078–2086.

64. Lobanenkov, V.V. and Zentner, G.E. (2018) Discovering a binary CTCF code with a little help from BORIS. Nucleus, 9, 33–41.

65. Owens, N., Papadopoulou, T., Festuccia, N., Tachtsidi, A., Gonzalez, I., Dubois, A., Vandormael-Pournin, S., Nora, E.P., Bruneau, B.G., Cohen-Tannoudji, M. et al. (2019) CTCF confers local nucleosome resiliency after DNA replication and during mitosis. bioRxiv, 563619.

66. Wiechens, N., Singh, V., Gkikopoulos, T., Schofield, P., Rocha, S. and Owen-Hughes, T. (2016) The Chromatin Remodelling Enzymes SNF2H and SNF2L Position Nucleosomes adjacent to CTCF and Other Transcription Factors. PLOS Genetics, 12, e1005940.

67. Barutcu, A.R., Lian, J.B., Stein, J.L., Stein, G.S. and Imbalzano, A.N. (2017) The connection between BRG1, CTCF and topoisomerases at TAD boundaries. Nucleus, 8, 150–155.

68. Ishihara, K., Oshimura, M. and Nakao, M. (2006) CTCF-Dependent Chromatin Insulator Is Linked to Epigenetic Remodeling. Molecular Cell, 23, 733–742.

69. Bornelov, S., Reynolds, N., Xenophontos, M., Gharbi, S., Johnstone, E., Floyd, R., Ralser, M., Signolet, J., Loos, R., Dietmann, S. et al. (2018) The Nucleosome Remodeling and Deacetylation Complex Modulates Chromatin Structure at Sites of Active Transcription to Fine-Tune Gene Expression. Mol Cell, 71, 56–72 e54.

70. Snyder, M.W., Kircher, M., Hill, A.J., Daza, R.M. and Shendure, J. (2016) Cell-free DNA Comprises an In Vivo Nucleosome Footprint that Informs Its Tissues-Of-Origin. Cell, 164, 57–68.

71. Hansen, A.S., Pustova, I., Cattoglio, C., Tjian, R. and Darzacq, X. (2017) CTCF and cohesin regulate chromatin loop stability with distinct dynamics. Elife, 6.

72. Barrington, C., Georgopoulou, D., Pezic, D., Varsally, W., Herrero, J. and Hadjur, S. (2019) Enhancer accessibility and CTCF occupancy underlie asymmetric TAD architecture and cell type specific genome topology. Nature communications, 10, 2908.

73. Ho, L., Jothi, R., Ronan, J.L., Cui, K., Zhao, K. and Crabtree, G.R. (2009) An embryonic stem cell chromatin remodeling complex, esBAF, is an essential component of the core pluripotency transcriptional network. Proc Natl Acad Sci U S A, 106, 5187–5191.

