## Supplementary Materials for "CTCF-dependent chromatin boundaries formed by asymmetric nucleosome arrays with decreased linker length" (from another version of this preprint: version 2, 2019-07-05)

[3] Wellcome Trust Vacation Student. Current address: Department of Biological and Medical Sciences, Oxford Brookes University, Headington Campus, Oxford, OX3 0BP, UK

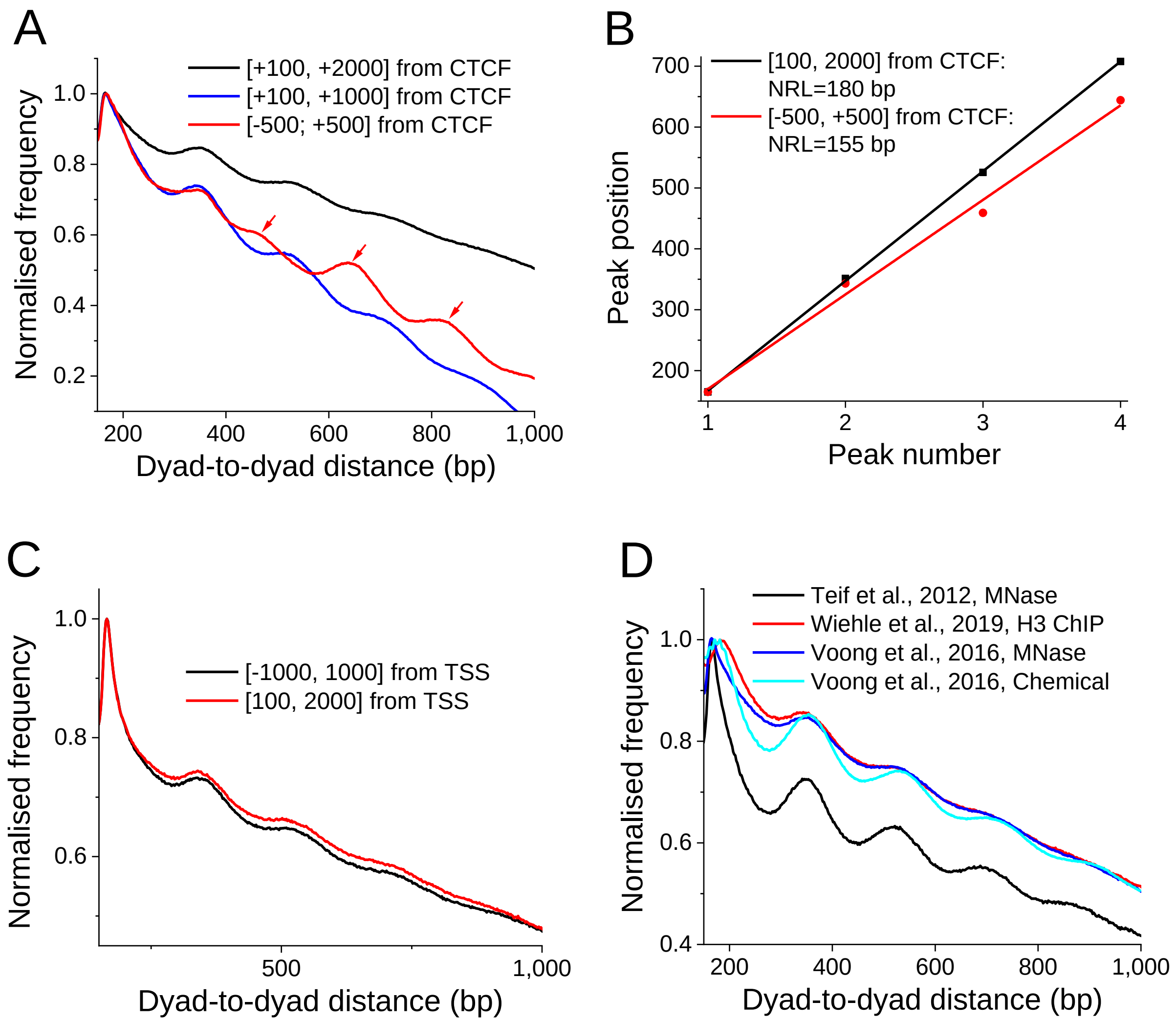


**Figure S1. The effect of region location near CTCF on the apparent NRL value.** A) Phasograms depicting the normalised frequency of nucleosome dyad-to-dyad distances calculated using NucTools for three different regions near CTCF sites: [100, 2000], [100, 1000] and [-500, 500]. Both [100, 2000] and [100, 1000] patterns oscillate with NRL=174. In the case of the [-500, 500] phasogram additional peaks (indicated by red arrows) appear which correspond to distances between nucleosomes on different sides of CTCF, thus resulting in an apparent NRL<160. B) NRLs calculated from the phasograms shown in panel (A). Region [-500, 500] is characterised by an unrealistically small NRL=155bp which is an artefact of the interference of two waves of distances between nucleosomes located on different sides from CTCF. C) Comparison of the phasograms showing the normalised frequency of nucleosome dyad-to-dyad distances for the regions [-1000, 1000] and [100, 2000] from TSS. The NRLs calculated based on these phasograms are not significantly different (172+/-1 bp vs. 168+/-3 bp respectively). D) Phasograms calculated for the region [100, 2000] near CTCF site for two different MNase-seq datasest from (1) and (2), MNase-assisted H3 ChIP-seq from (3) and chemical mapping from (2).

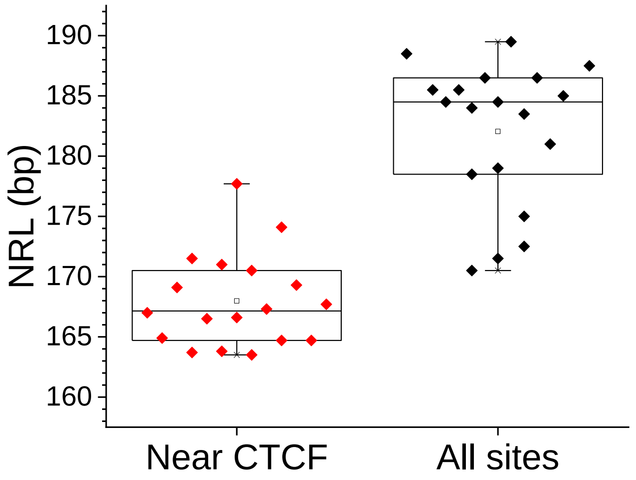


**Figure S2.** NRLs calculated near binding sites of 18 stemness-related chromatin proteins in ESCs in the region [250, 1000] from the TF sites. The same TF datasets as in Figure 1D are used. Left: TF binding sites in the vicinity of CTCF; right: all TF binding sites irrespective of their distance from CTCT.

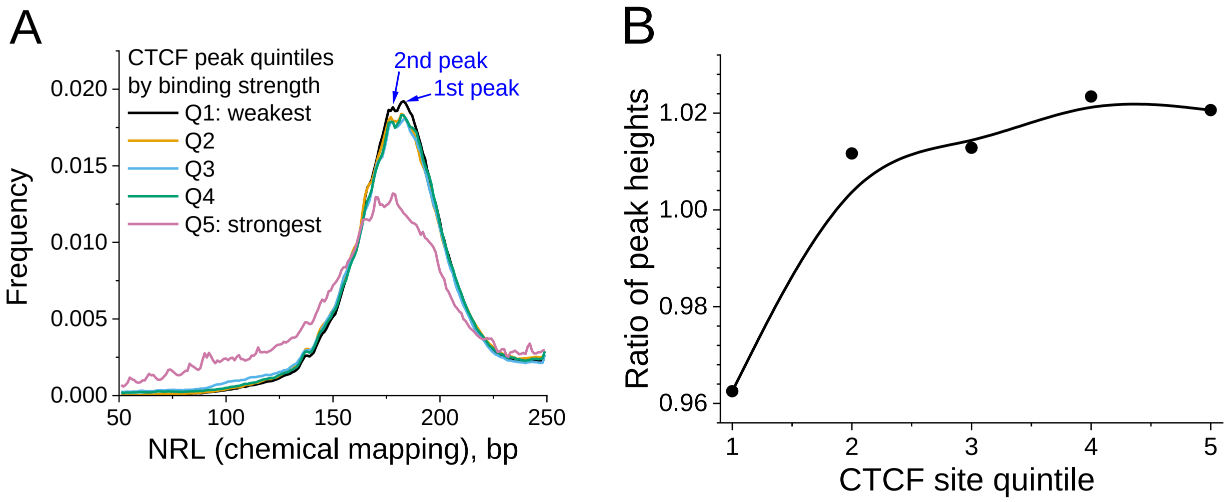


**Figure S3.** A) A histogram of nucleosome dyad-to-dyad distances determined using chemical mapping for different CTCF quintiles defined based on CTCF binding strength determined by the height of ChIP-seq CTCF peaks. Note that the dyad-to-dyad distances determined using chemical mapping cannot be directly compared to MNase-seq based NRLs due to an inherent bias of the chemical mapping experimental setup that we discussed previously (4). B) The ratio between heights of $2^{nd}$ peak and $1^{st}$ peak of the distribution of lengths of chemical mapping-based dyad-to-dyad distances shown in panel (A) as a function of the CTCF site quintile based on the heights of CTCF ChIP-seq peaks.

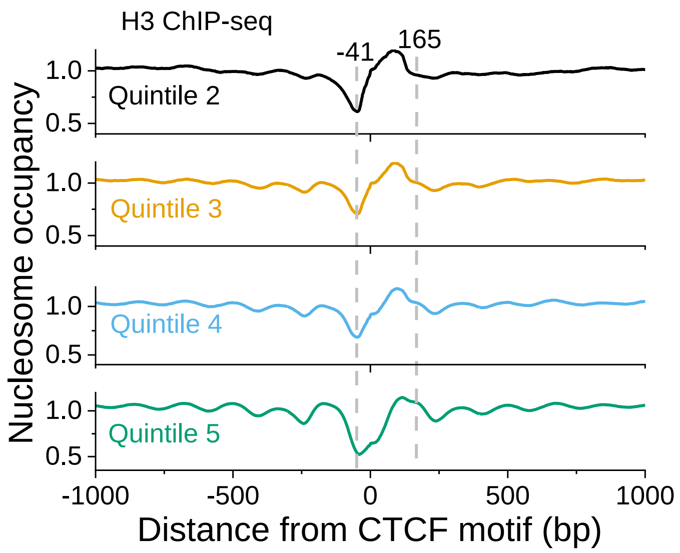


**Figure S4.** Aggregate nucleosome occupancy profiles around directional CTCF as in Figure 4, calculated using MNase-assisted histone H3 ChIP-seq signal from (Wiehle et al., 2019).

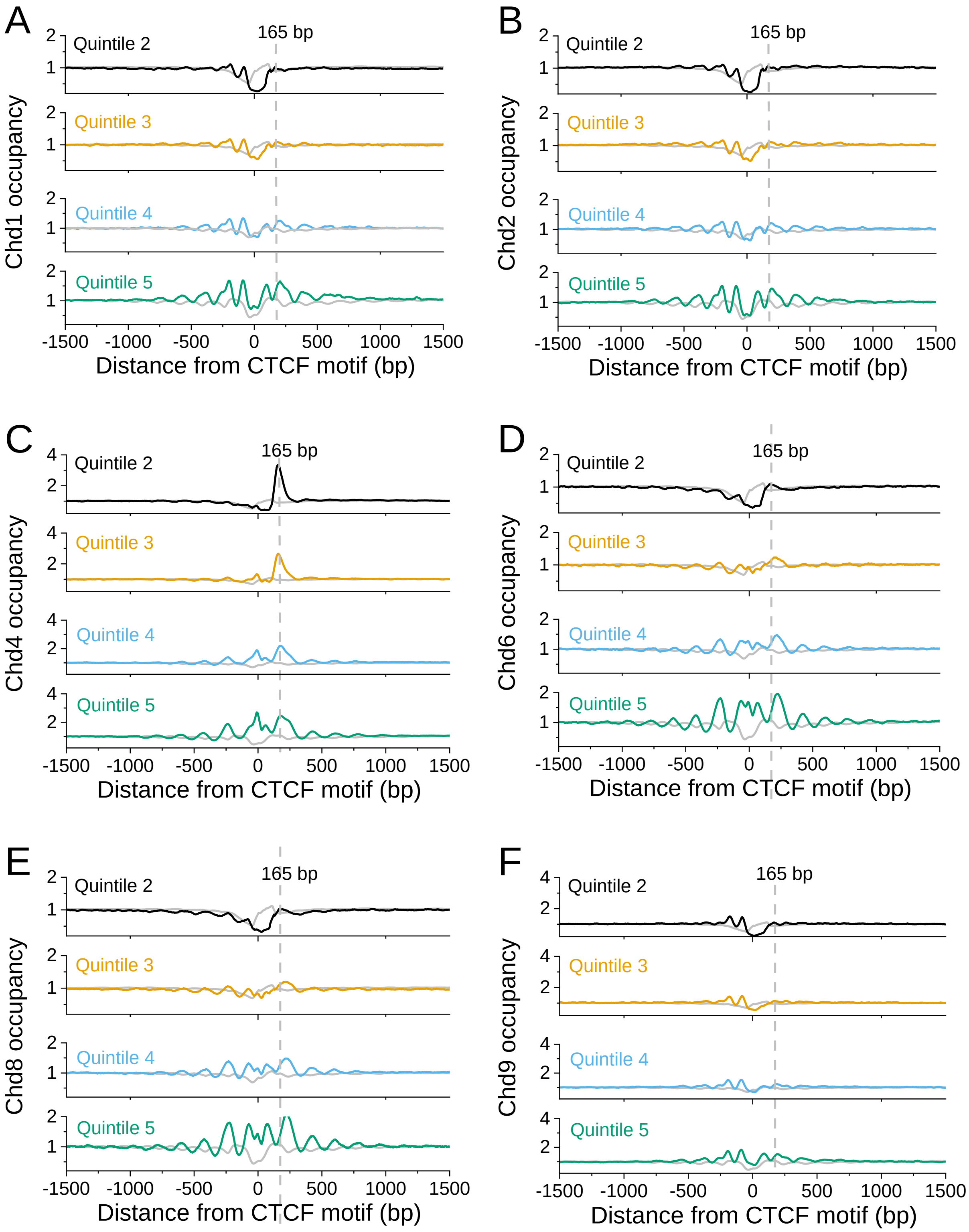

A
B
C
D
E
F
Quintile 2
Quintile 3
Quintile 4
Quintile 5
165 bp
Chd1 occupancy
Chd2 occupancy
Chd4 occupancy
Chd6 occupancy
Chd8 occupancy
Chd9 occupancy
Distance from CTCF motif (bp)
-1500
-1000
-500
0
500
1000
1500

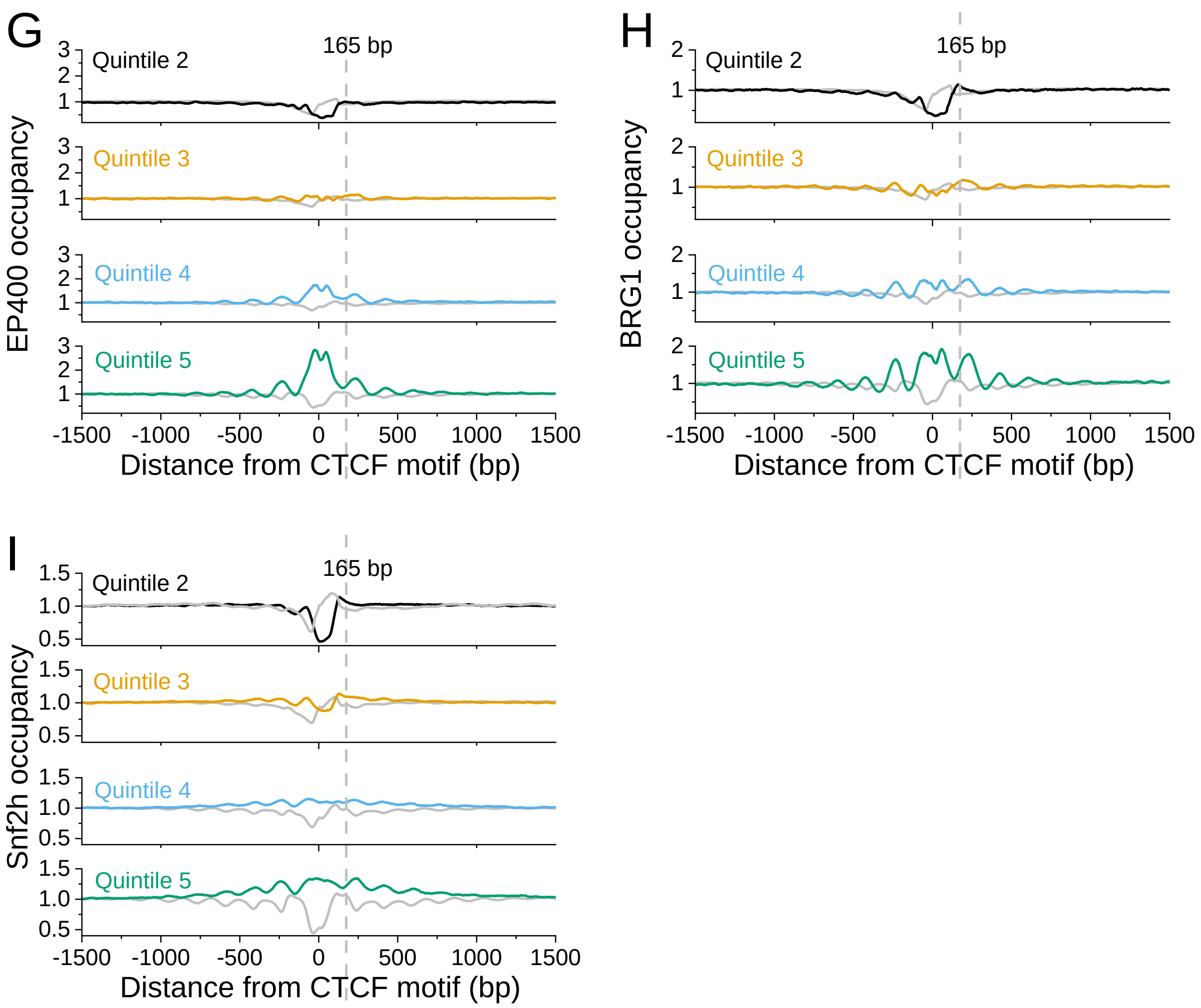


**Figure S5. Aggregate profiles of the occupancy of chromatin remodellers near directional CTCF motifs in ESCs.** Aggregate occupancy profiles for nine chromatin remodellers around all predicted CTCF motifs in the mouse genome show that only Chd4 has a CTCF-dependent peak at 165 bp (note a different scale for Chd4). Remodeller profiles are aligned around CTCF motifs split into quintiles with increasing CTCF binding strength as follows: black – 2$^{nd}$ quintile; orange – 3$^{rd}$ quintile; blue – 4$^{th}$ quintile; green – 5$^{th}$ quintile. Grey solid lines show the corresponding nucleosome occupancy based on MNase-seq as in Figure 5.

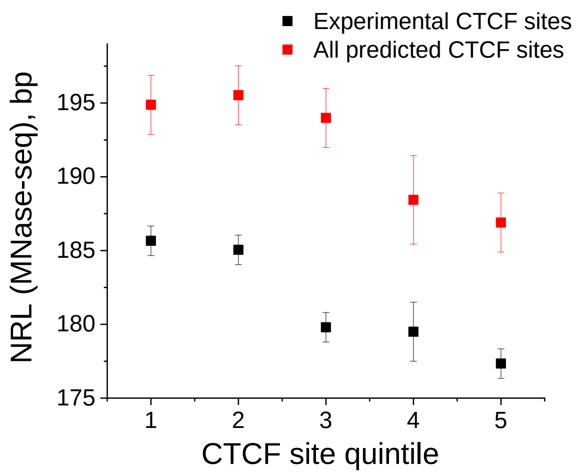


**Figure S6. NRL in the region [300, 2000] near CTCF as a function of CTCF binding strength.** The effect of NRL decrease with increase of CTCF binding strength remains even after excluding the CTCF-dependent nucleosome at +165 bp and performing NRL calculation for the region [300, 2000] downstream of CTCF sites (to the right from CTCF using plus strand coordinates).

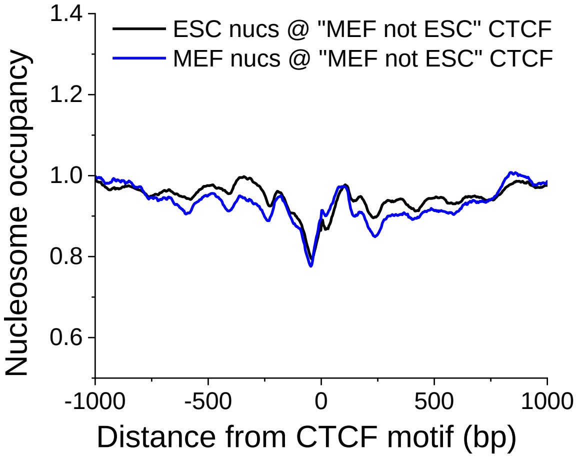


**Figure S7. Effects of CTCF-dependent boundary directionality in stem cell differentiation.** Nucleosome occupancy in ESCs (black) and MEFs (blue) around CTCF sites "MEF not ESC" that are present in MEFs but not in ESCs, calculated taking into account CTCF motif directionality.

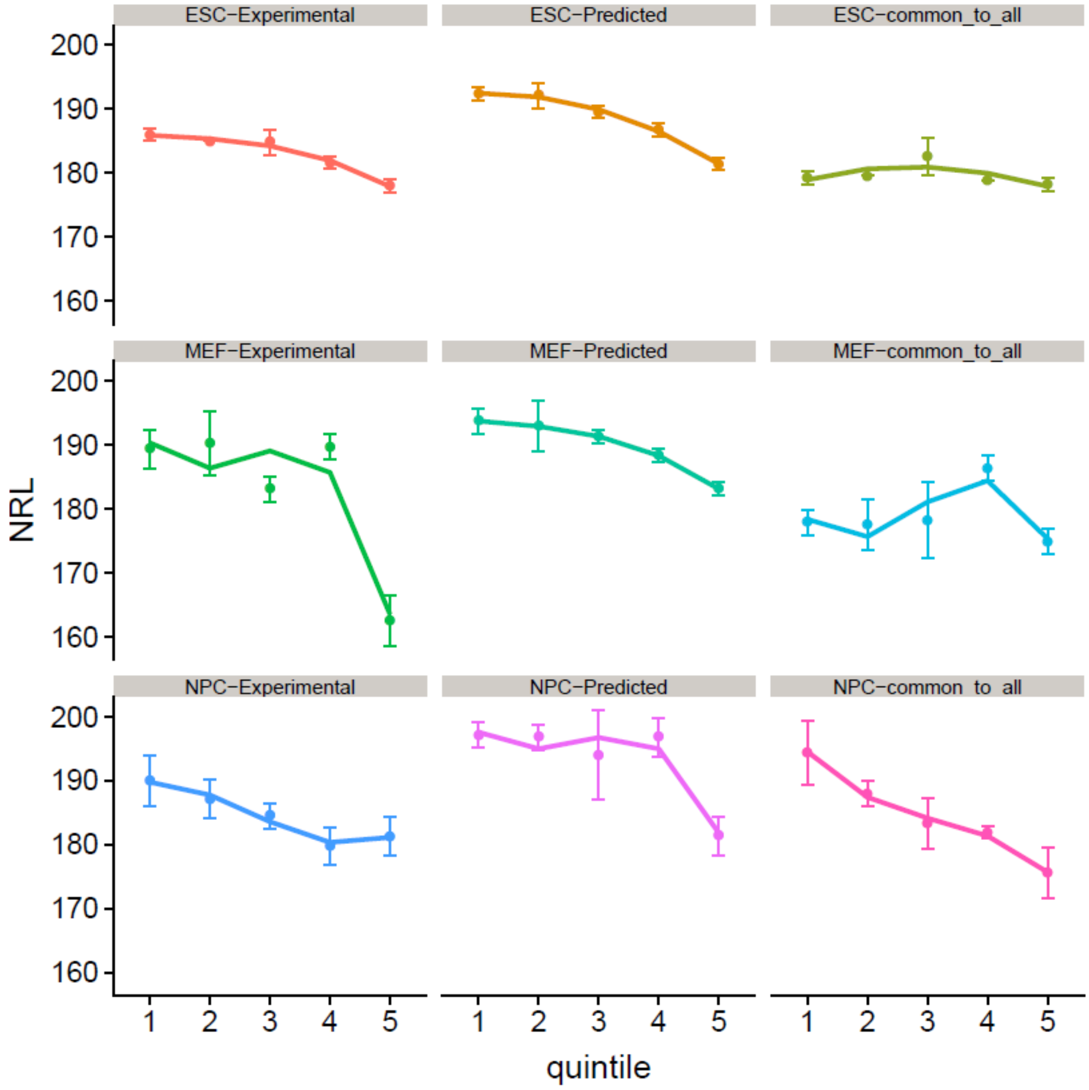


**Figure S8. Effect of ESC differentiation on NRL.** NRL values are calculated as a function of the CTCF site quintile. Top row: NRLs calculated based on the MNase-seq dataset in ESCs from Teif et al., 2012 for all experimental CTCF sites in ESCs determined in Shen et al, 2012 (left), all computationally predicted CTCF sites (middle) and common CTCF sites that have been determined experimentally in each of ESCs, NPCs and MEFs (right). Middle row: NRLs calculated based on the MNase-seq dataset in MEFs from Teif et al., 2012 for all experimental CTCF sites in MEF determined in Shen et al, 2012 (left), all computationally predicted CTCF sites (middle) and common CTCF sites that have been determined experimentally in each of ESCs, NPCs and MEFs (right). Bottom row: NRLs calculated based on the MNase-seq dataset in NPCs from Teif et al., 2012 for all experimental CTCF sites in NPCs determined in Bonev et al., 2017 (left), all computationally predicted CTCF sites (middle) and common CTCF sites that have been determined experimentally in each of ESCs, NPCs and MEFs (right).

**Supplemental Table ST1. Summary of experimental datasets used in this study**

| Name | Accession # | Reference |
|---|---|---|
| Nanog in ESC | GSM3123484 | (5) |
| Oct4 in ESC | GSM2417142 | (6) |
| Sox2 in ESC | GSM2417143 | (6) |
| Klf4 in ESC | GSM2417144 | (6) |
| cMyc in ESC | GSM2417145 | (6) |
| Esrrb in ESC | GSM2561449 | (7) |
| GATA1 in ESC | GSM453997 | (8) |
| Ep300 in ESC | ENCSR000CCD GSM918750 | (9) |
| Tal1 in ESC | ENCSR000DIN GSM923579 | (9) |
| Zbtb2 in ESC | GSM2716083 | (10) |
| ZNF384 in ESC | ENCSR000ERV GSM1003807 | (9) |
| CTCF in ESC and MEF | ENCSR000CCB GSM918748 | (9) |
| STAT3 in ESC | GSM288353 | (9) |
| GATA2 in ESC | ENCSR000DIE GSM923587 | (9) |
| cJun in ESC | GSM1587320 | (11) |
| Max in ESC | GSM1171650 | (12) |
| Arid3a in ESC | GSM1370509 | (13) |
| Sox17 in ESC | GSM1059856 | (14) |
| CpG islands | Obtained from authors' web site | (15) |
| BRG1 in ESC | GSM359413 | (16) |
| Snf2h in ESC | GSE80049 | (17) |
| Nucleosome Chemical mapping in ESC | GSE82127 | (2) |
| MNase-seq in ESC (Voong et al., 2016) | GSM2183911 | (2) |
| MNase-seq (Teif et al, 2012, 2014) | GSE40896 | (1,18) |
| MNase-H3-ChIP-seq | GSE114599 | (3) |
| BRG1 in ESC | GSE64825 | (19) |
| Chd1 in ESC | GSE64825 | (19) |
| Chd2 in ESC | GSE64825 | (19) |
| Chd4 in ESC | GSE64825 | (19) |

| | | |
|---|---|---|
| Chd6 in ESC | GSE64825 | (19) |
| Chd8 in ESC | GSE64825 | (19) |
| Chd9 in ESC | GSE64825 | (19) |
| EP400 in ESC | GSE64825 | (19) |
| Chromatin loops and TADs in ESC | BED files provided by the authors | (20) |
| Promoters | RefSeq | (21) |
| Enhancers | http://fantom.gsc.riken.jp (all permissive enhancers) | (22) |
| CTCF in ESC and MEF | GSE114599 | (3) |
| CTCF in ESC and MEF | GSE27944 | (23) |
| CTCF in ESC and NPC | GSE96107 | (20) |

Supplementary references


1. Teif, V.B., Vainshtein, Y., Caudron-Herger, M., Mallm, J.P., Marth, C., Höfer, T. and Rippe, K. (2012) Genome-wide nucleosome positioning during embryonic stem cell development. *Nat Struct Mol Biol*, **19**, 1185-1192.
2. Voong, L.N., Xi, L., Sebeson, A.C., Xiong, B., Wang, J.P. and Wang, X. (2016) Insights into Nucleosome Organization in Mouse Embryonic Stem Cells through Chemical Mapping. *Cell*, **167**, 1555-1570 e1515.
3. Wiehle, L., Thorn, G.J., Raddatz, G., Clarkson, C.T., Rippe, K., Lyko, F., Breiling, A. and Teif, V.B. (2019) DNA (de)methylation in embryonic stem cells controls CTCF-dependent chromatin boundaries. *Genome Res*, **29**, 750-761.
4. Vainshtein, Y., Rippe, K. and Teif, V.B. (2017) NucTools: analysis of chromatin feature occupancy profiles from high-throughput sequencing data. *BMC Genomics*, **18**, 158.
5. Kim, K.Y., Tanaka, Y., Su, J., Cakir, B., Xiang, Y., Patterson, B., Ding, J., Jung, Y.W., Kim, J.H., Hysolli, E. *et al.* (2018) Uhrf1 regulates active transcriptional marks at bivalent domains in pluripotent stem cells through Setd1a. *Nature communications*, **9**, 2583.
6. Chronis, C., Fiziev, P., Papp, B., Butz, S., Bonora, G., Sabri, S., Ernst, J. and Plath, K. (2017) Cooperative Binding of Transcription Factors Orchestrates Reprogramming. *Cell*, **168**, 442-459 e420.
7. Xie, L., Torigoe, S.E., Xiao, J., Mai, D.H., Li, L., Davis, F.P., Dong, P., Marie-Nelly, H., Grimm, J., Lavis, L. *et al.* (2017) A dynamic interplay of enhancer elements regulates Klf4 expression in naive pluripotency. *Genes Dev*, **31**, 1795-1808.
8. Chen, X., Xu, H., Yuan, P., Fang, F., Huss, M., Vega, V.B., Wong, E., Orlov, Y.L., Zhang, W., Jiang, J. *et al.* (2008) Integration of external signaling pathways with the core transcriptional network in embryonic stem cells. *Cell*, **133**, 1106-1117.
9. Shen, Y., Yue, F., McCleary, D.F., Ye, Z., Edsall, L., Kuan, S., Wagner, U., Dixon, J., Lee, L., Lobanenkov, V.V. *et al.* (2012) A map of the cis-regulatory sequences in the mouse genome. *Nature*, **488**, 116-120.

10. Karemaker, I.D. and Vermeulen, M. (2018) ZBTB2 reads unmethylated CpG island promoters and regulates embryonic stem cell differentiation. *EMBO Rep*, **19**.
11. Liu, J., Han, Q., Peng, T., Peng, M., Wei, B., Li, D., Wang, X., Yu, S., Yang, J., Cao, S. *et al.* (2015) The oncogene c-Jun impedes somatic cell reprogramming. *Nat Cell Biol*, **17**, 856-867.
12. Krepelova, A., Neri, F., Maldotti, M., Rapelli, S. and Oliviero, S. (2014) Myc and max genome-wide binding sites analysis links the Myc regulatory network with the polycomb and the core pluripotency networks in mouse embryonic stem cells. *PLoS One*, **9**, e88933.
13. Rhee, C., Lee, B.-K., Beck, S., Anjum, A., Cook, K.R., Popowski, M., Tucker, H.O. and Kim, J. (2014) Arid3a is essential to execution of the first cell fate decision via direct embryonic and extraembryonic transcriptional regulation. *Genes & Development*, **28**, 2219-2232.
14. Aksoy, I., Jauch, R., Chen, J., Dyla, M., Divakar, U., Bogu, G.K., Teo, R., Leng Ng, C.K., Herath, W., Lili, S. *et al.* (2013) Oct4 switches partnering from Sox2 to Sox17 to reinterpret the enhancer code and specify endoderm. *EMBO J*, **32**, 938-953.
15. Irizarry, R.A., Wu, H. and Feinberg, A.P. (2009) A species-generalized probabilistic model-based definition of CpG islands. *Mamm Genome*, **20**, 674-680.
16. Ho, L., Jothi, R., Ronan, J.L., Cui, K., Zhao, K. and Crabtree, G.R. (2009) An embryonic stem cell chromatin remodeling complex, esBAF, is an essential component of the core pluripotency transcriptional network. *Proc Natl Acad Sci U S A*, **106**, 5187-5191.
17. Local, A., Huang, H., Albuquerque, C.P., Singh, N., Lee, A.Y., Wang, W., Wang, C., Hsia, J.E., Shiau, A.K., Ge, K. *et al.* (2018) Identification of H3K4me1-associated proteins at mammalian enhancers. *Nat Genet*, **50**, 73-82.
18. Teif, V.B., Beshnova, D.A., Vainshtein, Y., Marth, C., Mallm, J.P., Höfer, T. and Rippe, K. (2014) Nucleosome repositioning links DNA (de)methylation and differential CTCF binding during stem cell development. *Genome Res*, **24**, 1285-1295.
19. de Dieuleveult, M., Yen, K., Hmitou, I., Depaux, A., Boussouar, F., Bou Dargham, D., Jounier, S., Humbertclaude, H., Ribierre, F., Baulard, C. *et al.* (2016) Genome-wide nucleosome specificity and function of chromatin remodellers in ES cells. *Nature*, **530**, 113-116.
20. Bonev, B., Mendelson Cohen, N., Szabo, Q., Fritsch, L., Papadopoulos, G.L., Lubling, Y., Xu, X., Lv, X., Hugnot, J.P., Tanay, A. *et al.* (2017) Multiscale 3D Genome Rewiring during Mouse Neural Development. *Cell*, **171**, 557-572 e524.
21. Pruitt, K.D., Brown, G.R., Hiatt, S.M., Thibaud-Nissen, F., Astashyn, A., Ermolaeva, O., Farrell, C.M., Hart, J., Landrum, M.J., McGarvey, K.M. *et al.* (2014) RefSeq: an update on mammalian reference sequences. *Nucleic Acids Res*, **42**, D756-763.
22. Lizio, M., Harshbarger, J., Shimoji, H., Severin, J., Kasukawa, T., Sahin, S., Abugessaisa, I., Fukuda, S., Hori, F., Ishikawa-Kato, S. *et al.* (2015) Gateways to the FANTOM5 promoter level mammalian expression atlas. *Genome Biol*, **16**, 22.
23. Martin, D., Pantoja, C., Fernandez Minan, A., Valdes-Quezada, C., Molto, E., Matesanz, F., Bogdanovic, O., de la Calle-Mustienes, E., Dominguez, O., Taher, L. *et al.* (2011) Genome-wide CTCF distribution in vertebrates defines equivalent sites that aid the identification of disease-associated genes. *Nat Struct Mol Biol*, **18**, 708-714.
